# Planarian behavioral screening is a useful invertebrate model for evaluating seizurogenic chemicals

**DOI:** 10.1101/2025.10.07.680963

**Authors:** Danielle Ireland, Evangeline Coffinas, Christina Rabeler, Eva-Maria S. Collins

## Abstract

Detecting adverse health effects of drugs and other chemicals early during chemical/drug development saves significant time and resources. Freshwater planarians are an emerging invertebrate model for rapid, cost-effective neurotoxicity screening. Because planarians exhibit seizure-like behavior when exposed to chemicals that cause seizures in mammals, such as N-methyl-D-aspartate (NMDA) and picrotoxin, they could be a useful first-tier model for seizure screening and thus reduce the need for slow and expensive mammalian tests. However, planarian seizure studies to date have been low-throughput and lacking the necessary standardization and automated analysis to make this model a viable screening solution. Here, we present results from medium-throughput behavioral testing conducted in 48-well plates using two popular models for planarian pharmacological and toxicological studies: *Dugesia japonica* and *Girardia dorotocephala*. Planarian behavior was scored using automated image analysis, measuring both translational behavior and body shape changes. We found that known seizurogenic compounds in mammals (NMDA, nicotine, picrotoxin, pilocarpine, and pentylenetetrazole (PTZ)) induced seizure-like behavior in both planarian species within 30 minutes of exposure. We also tested three pesticides (parathion, carbaryl, and permethrin). Parathion and carbaryl, but not permethrin, caused planarian seizure-like activity. While the planarian species responded similarly to most compounds, some compounds showed potency differences of 10-100-fold (pilocarpine and nicotine, respectively). *G. dorotocephala* planarians were generally more sensitive, but *D. japonica* planarians displayed more reproducible behaviors. By standardizing both experimental approach and analysis methods and making them available, this work can serve as a framework for future testing of chemicals for seizurogenic potential in planarians.

## 1. Introduction

Drugs that target the central nervous system (CNS) or non-CNS drugs that can cross the blood brain barrier may cause adverse outcomes on brain function, such as the induction of seizures. In fact, a recent study (Behl et al., 2025) reports that failure during seizure liability testing accounts for 67% of failures of CNS-drugs during pre-clinical drug development, based on survey results by the Safety Pharmacology Society in 2015 (Authier et al., 2016). Seizures can be induced through different molecular initiation events (Behl et al., 2025; Jett, 2012; Tukker and Westerink, 2021). For example, the well-known seizurogenic chemicals pentylenetetrazole (PTZ) and picrotoxin (PTX) cause seizures by inhibiting the GABA_A_ receptor (Squires et al., 1984) while acute toxic exposure to organophosphorus nerve gases and pesticides causes seizures due to inhibition of acetylcholinesterase (AChE) (Kozhemyakin et al., 2010). Thus, seizure liability testing is an important step during pre-clinical drug development for drugs that enter the CNS or possess structural similarity to known seizurogenic compounds (Behl et al., 2025; Easter et al., 2009; Roberts et al., 2021).

Detection of potential seizurogenic compounds has traditionally relied on mice and rat *in vivo* and *ex vivo* models during late stage preclinical studies (Easter et al., 2009; Tukker and Westerink, 2021). However, there is increasing awareness that testing in rodents is too expensive and slow to keep up with the demands for chemical screening and may be limited in its predictive capabilities for human health (Behl et al., 2025; Smirnova et al., 2014; Van Norman, 2019). Therefore, multiple *in silico* and *in vitro* assays have been developed in recent years to fill the need to more quickly and efficiently identify seizure liability (Behl et al., 2025; Belair et al., 2025; Easter et al., 2009; Grainger et al., 2018; Tukker and Westerink, 2021). Specifically, micro-electrode arrays (MEAs), particularly those utilizing human induced pluripotent stem cell derived neuronal /astrocyte co-cultures, have emerged as a promising new approach method (Belair et al., 2025; Grainger et al., 2018; Tukker and Westerink, 2021). In these assays, whole network function can be quantified and specific changes in neuronal activity have been shown to be indicative of *in vivo* seizures (Ishii et al., 2017). However, cell systems, including 3D cultures (Pelkonen et al., 2020), lack the complexity and metabolic capacity necessary to fully evaluate the effects of chemicals on brain function in a living organism (Hogberg and Smirnova, 2022). Small organismal models help overcome these challenges as they contain the entire complexity of the nervous system in its original context and have metabolic capacity, albeit with species-dependent differences, and thus can bridge between *in vitro* and mammalian models (Collins et al., 2024). Developing zebrafish allow for rapid and cost-effective screening in an intact organism and therefore have become a popular complementary test system, especially for rapid screening of anti-seizure compounds (Baraban et al., 2005; McGraw and Poduri, 2025; Milder et al., 2022; Whyte-Fagundes et al., 2025).

No model can 100% predict human *in vivo* effects and therefore modern toxicology aims to leverage the evidence gained from multiple complementary models via integrated approaches to hazard and risk assessment (Hogberg et al., 2022; Jaworska et al., 2016; Rockley et al., 2023). Freshwater planarians are invertebrates that have been utilized for pharmacological studies for over a century (Buttarelli et al., 2008; Ireland and Collins, 2022; Raffa, 2008). Additionally, planarians have recently gained recognition for their amenability to rapid behavioral testing of toxicants and neuroactive drugs (Ireland et al., 2025a; Ireland and Collins, 2022, 2023). Planarians are sensitive to chemical exposure and respond with stereotypical behavioral responses that can be measured (reviewed in (Buttarelli et al., 2008; Hagstrom et al., 2016; Raffa, 2008)). Chemically-induced behaviors have been qualitatively described in the planarian literature, but with the exception of “scrunching”, a well-characterized planarian behavior that shows oscillatory body shape changes (Cochet-Escartin et al., 2015; Sabry et al., 2019), chemically-induced behaviors remain to be quantitatively defined (Reho et al., 2022). Because planarians live in water, chemical exposure is simple and behavioral experiments can be conducted easily and inexpensively (Ireland and Collins, 2022). Planarians share key neurotransmitter systems with humans, including GABA, glutamate, serotonin, dopamine, and acetylcholine (Ross et al., 2017), thus making them a relevant neurobiology model. Different species have been used for chemical exposure experiments for different reasons, such as commercial availability, size, and more recently genome availability and amenability to high-throughput studies in multi-well plates (Ireland and Collins, 2023; Wu and Li, 2018).

Studies focused specifically on seizure-like behavior in planarians (Figure 1) have primarily used the commercially available *Girardia* (formerly *Dugesia*) *dorotocephala* (GD) and *Girardia* (formerly *Dugesia) tigrina* (GT) species and found that these species are sensitive to known seizurogenic compounds such as N-methyl-D-aspartate (NMDA) (Rawls et al., 2009), nicotine (Bach et al., 2016; Pagán et al., 2015; Rawls et al., 2011), and pilocarpine (Miller et al., 2025). Because it has not yet been shown that the behavioral phenotypes induced by these compounds are due to seizures through EEG or other means to visualize neuronal activity, we will refer to them as ‘planarian seizure-like activity’ (pSLA), a term originally introduced by (Rawls et al., 2009) and defined as “asynchronous paroxysms (C-shape, twitching behavior)”. Other researchers have also referred to pSLA as planarian seizure-like motility (pSLM) (Pagán et al., 2013), defined as “vigorous writhing and bending of the body” (Bach et al., 2016). A clear definition of what these terms encapsulate does not exist but they seem to denote rapid shape changes – each lasting approximately 1 second (Rawls et al., 2009) - that include C-shape, writhing, head flailing, dorsal oscillation, corkscrew, among others. Some of these shapes are well-defined (e.g., corkscrew) and examples have previously been reviewed (Hagstrom et al., 2016; Wu and Li, 2018), while others are ambiguous, e.g., “dorsal oscillations” (Miller et al., 2025).

**Figure 1.**
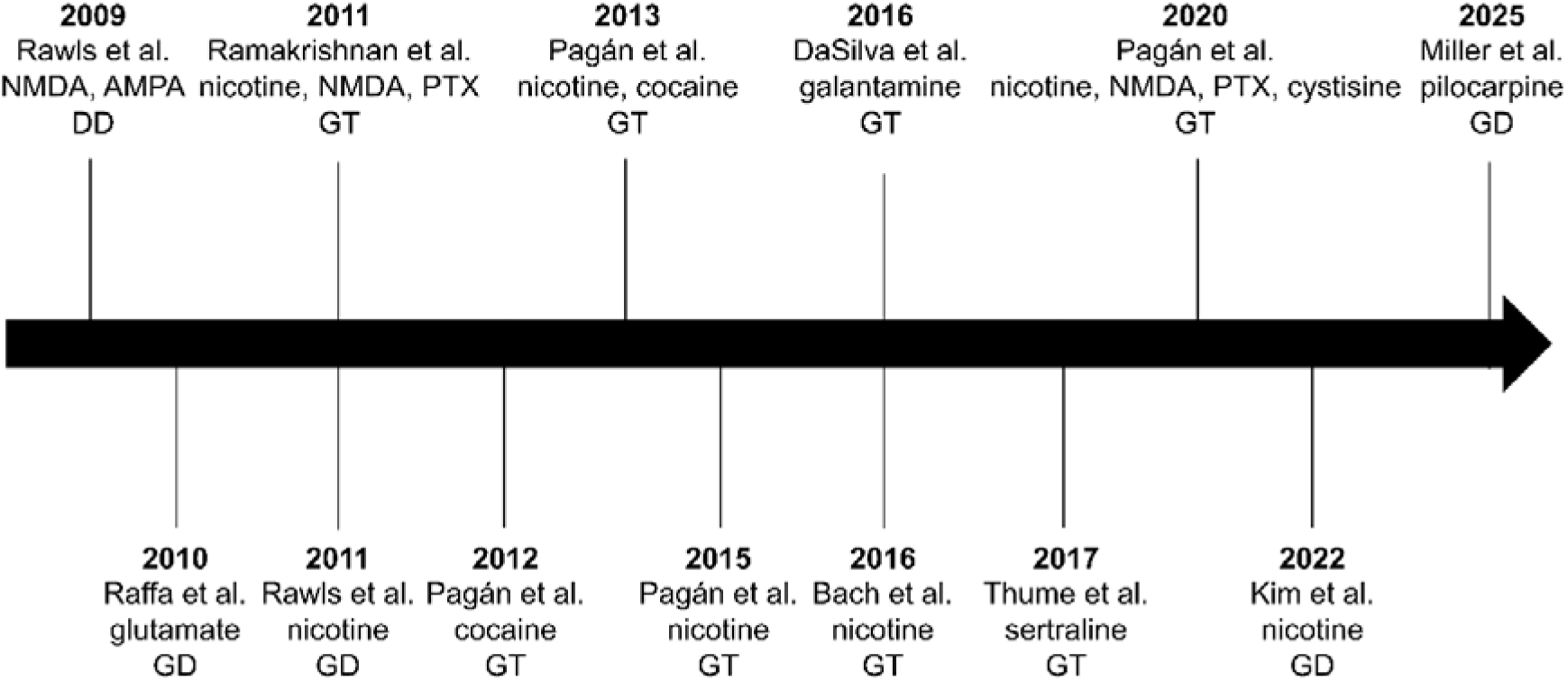
Overview of planarian studies on seizurogenic chemicals in the last 25 years (2000-2025). Tested seizurogenic compounds are listed. Some of the studies also tested other compounds, including therapeutics to treat seizures, which are not listed here. Abbreviations indicate the species tested: GD – *Girardia dorotocephala*; GT – Girardia *tigrina*.

While object tracking has been employed to characterize the spatial behavior of planarians exposed to seizurogenic chemicals (Miller et al., 2025), pSLA thus far has been scored manually, by counting the number of abnormal behavioral changes during a 5-10 minute recording period. Manual pSLA scoring relies on qualitative criteria about shape changes, which may differ between laboratories and even researchers. Moreover, behavioral data is intrinsically noisy due to the biological variability of organisms; thus, standardization of methods to minimize variability and automated pSLA analysis is indispensable to overcome these limitations and guarantee reproducibility and transferability of results.

Here, we present a medium-throughput method for screening chemicals for seizurogenic potential in two planarian species that are popular models in toxicology and pharmacology research: *Dugesia japonica* (DJ) and GD planarians. Screening is performed in 48-well plates and scoring of pSLA is achieved for all 48 planarians simultaneously via automated image analysis using a Motility Index (MI) metric that uses a standard image subtraction algorithm. The MI method is similar to methods routinely used to evaluate zebrafish behavior (Bruni et al., 2016), including specifically for seizures (e.g., (Afrikanova et al., 2013; Whyte-Fagundes et al., 2025)). The 48-well plate format and automated analysis greatly increases the throughput from previous planarian studies, which utilized exposure in petri dishes (e.g., (Bach et al., 2016; Rawls et al., 2009)) or 12-well plates (Miller et al., 2025) and used primarily manual pSLA scoring of individual planarians. Manual scoring of 48 worms recorded for 30 minutes would take 24 hours compared to ∼15 minutes of run time of the automated computational analysis we present here. Moreover, automated analysis allows for a standardized, quantitative definition of pSLA that can be used across laboratories.

Using this automated analysis, we quantified the behavior of DJ and GD planarians to 9 test compounds. Four of the compounds have previously been shown to induce pSLA in some planarian species (NMDA, nicotine, picrotoxin (PTX), pilocarpine; Figure 1) and were confirmed as pSLA inducers by our automated analysis, which quantitatively defined pSLA based on species-specific thresholds in MI scores derived from qualitative scoring of NMDA-induced pSLA. PTZ, which is used to create rodent and zebrafish seizure models (Baraban et al., 2005; Dhir, 2012; Gawel et al., 2020; Milder et al., 2022; Shimada and Yamagata, 2018; Yang et al., 2023), also induced pSLA in both planarian species. We also evaluated allyl isothiocyanate (AITC) as a control compound that induces scrunching (Cochet-Escartin et al., 2015; Sabry et al., 2019) and show that our analysis can distinguish it from pSLA. The other test compounds were three pesticides with seizurogenic potential in mammals, namely two acetylcholinesterase inhibitors (parathion and carbaryl) and the pyrethroid permethrin (Jett, 2012). Permethrin, tested to the solubility limit, did not cause effects but parathion and carbaryl induced pSLA, thus demonstrating that pSLA can be triggered through different pathways in planarians, similar to in mammals.

## 2. Methods

### 2.1. Planarian species and maintenance

The freshwater planarian species DJ and GD were used in this study. DJ planarians have been maintained in the laboratory for > 15 years on organic liver (beef and chicken). GD planarians were obtained commercially from Carolina Biologicals (Cat #132954) and then maintained on frozen bloodworms (San Francisco Bay Brand, #88055) in the laboratory. All species were maintained in planarian water (0.21 g/L Instant Ocean salts [Spectrum Brands, Blacksburg, VA, USA], 0.83 mM MgSO_4_, 0.9 mM CaCl_2_, 0.04 mM KHCO_3_, 0.9mM NaHCO_3_), fed 1-2x per week and cleaned on the day of feeding and 2 days following feeding. Unless otherwise noted, all experiments used intact, adult planarians which had been fasted 4-9 days. Note that planarians can live for months without any food (McConnell, 1967) and we did not observe differences in response to the chemicals depending on the fasting state (Supplemental Figure S1). To determine if a brain is required for pSLA (section 3.6), planarians of similar size to intact ones were amputated 4 hours prior to chemical exposure with an ethanol-sterilized razor or scalpel blade between the auricles and pharynx.

### 2.2. Chemicals and preparation

Chemicals used in this study are listed in Table 1. For chemicals that had previously been tested in planarian seizure studies, test concentrations were guided by previous literature (Table 2), testing a range of concentrations around those shown to previously induce pSLA. For chemicals which had not previously been tested acutely in planarians, initial test concentration ranges were chosen based on either concentrations previously used in zebrafish to induce seizures (Afrikanova et al., 2013; Baraban et al., 2005) or concentrations which induced chronic effects in planarians (Ireland et al., 2022b; Zhang et al., 2019). The reported concentrations were refined by preliminary experiments to identify a concentration range spanning no effects to induction of pSLA or the solubility limit of the chemical, and thus were often shifted for the two planarian species. PTX was only tested at one concentration as this was the only concentration we found induced pSLA and higher concentrations were limited by solubility. Depending on water solubility, chemical stock solutions were prepared either in planarian water or in 100% dimethyl sulfoxide (DMSO) in a fume hood (Table 1). For all chemicals prepared in DMSO, the final DMSO concentration in the 1x solution for all concentrations was 0.5% (v/v) and 0.5% (v/v) DMSO was used as the solvent control for these chemicals. Because DMSO alone can affect planarian behavior (Pagán et al., 2006), we tested DMSO up to 5% for acute behavioral effects. No overt qualitative behavioral phenotypes or pSLA were found for DMSO at concentrations <4% in DJ or <5% in GD (Supplemental Figure S2). Chemicals prepared in planarian water were used the same day, while stocks made in DMSO were aliquoted and stored at −20C or −80C until use. Frozen stocks were thawed a maximum of 3 times (Kozikowski et al., 2003). Depending on chemical solubility, chemicals were added as 1x, 2x or 10x solutions to the planarians in the wells (Table 1).

**Table 1.** Overview of chemicals and concentrations used.

| Chemical | CAS # | Tested concentrations<sup>1</sup> | Vendor | Purity (%) |
| --- | --- | --- | --- | --- |
| allyl isothiocyanate (AITC)<sup>2</sup> | 57-06-7 | 25 µM (GD);<br>100 µM (DJ) | Sigma-Aldrich | 95% |
| carbaryl<sup>3</sup> | 63-25-2 | 10-75 µM | Sigma-Aldrich | ≥98.0 % |
| dimethyl sulfoxide (DMSO)<sup>2</sup> | 67-68-5 | 0.1%-5% (GD);<br>0.5-5% (DJ) | Sigma-Aldrich | ≥99.7% |
| N-methyl-D-aspartate (NMDA)<sup>4</sup> | 6384-92-5 | 3 mM (GD);<br>15 mM (DJ) | Tocris<br>Bioscience | ≥99% (HPLC) |
| (-)-nicotine<sup>2</sup> | 54-11-5 | 10-50 µM (GD);<br>50-1000 µM (DJ) | Sigma-Aldrich | ≥99% (GC) |
| parathion<sup>3</sup> | 56-38-2 | 10-50 µM (GD);<br>10-75 µM<sup>6</sup> (DJ) | Sigma-Aldrich;<br>ChemService | ≥98.0% (HPLC);<br>98.8% (GC/FID) |
| pentylentetrazole (PTZ)<sup>5</sup> | 54-95-5 | 5- 20 mM | Cayman<br>Chemical | ≥98% |
| permethrin (mixture of cis and trans isomers)<sup>3</sup> | 52645-53-1 | 100 µM; 1 mM<sup>6</sup> | Sigma-Aldrich | >97%<br>(29.8% cis,<br>67.8% trans<br>isomers) |
| picrotoxin,<br>*Anamirta cocculin* (PTX)<sup>4</sup> | 124-87-8 | 5 mM<sup>6</sup> | Sigma-Aldrich | ≥98% (mixture<br>of picrotoxin and<br>picrotin, HPLC) |
| (+)-pilocarpine hydrochloride<sup>5</sup> | 54-71-7 | 2-6 mM (GD); 6-100 mM (DJ) | Cayman<br>Chemical | ≥98% |

<sup>1</sup>if the same concentration or concentration range was used in both species, there is no species reference; else, it is indicated in parentheses which species was tested at this concentration.
<sup>2</sup> indicates highest concentration stock was 10x of the nominal, prepared in planarian water.
<sup>3</sup> indicates highest concentration stock was 200x of the nominal, prepared in DMSO and diluted to an intermediate 2x or 10x.
<sup>4</sup> indicates highest concentration stock was 1x of the nominal, prepared in planarian water.
<sup>5</sup> indicates highest concentration stock was 2x of the nominal, prepared in planarian water.
<sup>6</sup> indicates solubility issues with the 1x stock in planarian water.

**Table 2.** Chemicals and concentrations reported to induce pSLA in planarians.

| Chemical | Species | Concentration range | References |
| --- | --- | --- | --- |
| Nicotine | GT; GD | 10-2000 μM (GT)<br>1-5 mM (GD) | (Bach et al., 2016; Pagán et al., 2015; Ramakrishnan and DeSaer, 2011; Rawls et al., 2011) |
| Pilocarpine | GD | 1,2,3,4,6 mM | (Miller et al., 2025) |
| PTX | GT | 0.01-5 mM | (Ramakrishnan and DeSaer, 2011) |

pH measurements were taken with an Apera PH60-MS pH Tester kit (Apera Instruments, Columbus, OH) after allowing the probe to stabilize >10 minutes. The pH of the highest tested concentration of each chemical is listed in Supplemental Table S1. As the tested concentrations of NMDA were found to have very low pH (3-3.6), we compared NMDA exposure to planarian water at the same pH. To obtain planarian water at acidic pH, dilute hydrochloric acid was added until the desired pH was obtained and pH was verified immediately before use.

### 2.3. Plate setup and acute recording

Planarians were loaded into the wells of a tissue culture treated 48-well plate (Genesee Scientific, El Cajon, CA), one planarian per well, using a p1000 pipettor with the pipette tip slightly trimmed with a clean razor blade to avoid damage to the planarians. The planarians were loaded in planarian water and an appropriately concentrated chemical stock was added such that the total volume of 1x solution in each well equalled 200 µL. Chemical addition was performed in the fume hood. The chemical stock was added using a multichannel pipettor (CAPP Aero 12-Channel Pipette, 5-50 µL), the plate was sealed using thermal film (Excel Scientific, Victorville, CA, USA), and behavior was recorded, with the recordings beginning 2-6 minutes after chemical addition, including the transfer time between the fume hood and the recording setup and starting the recording. Four conditions (3 test conditions and 1 vehicle control) were tested per plate, with each condition comprising two columns of wells, with n=12 planarians per condition (chemical concentration and worm type) (Figure 2A). Separate plates were tested for each planarian species. If any chemical in the plate was prepared in DMSO, the vehicle control consisted of 0.5% (v/v) DMSO, else it contained untreated planarian water. As needed due to loss of worms crawling out of the wells or for cases with high data dispersion, independent replicates were performed to increase the sample size. The sample sizes for all experiments are listed in Supplemental Table S3.

**Figure 2.**
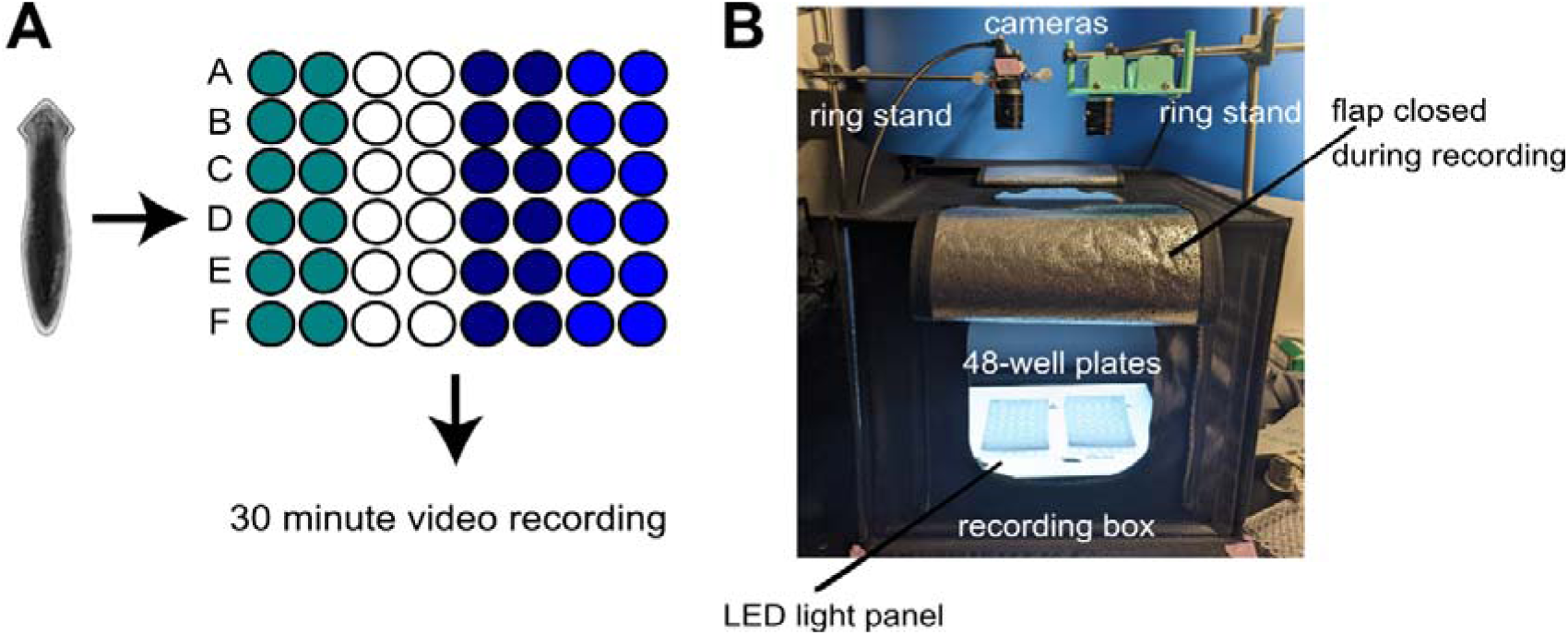
Schematic of 48-well plate and behavioral imaging setup. A) Testing was performed in 48-well plates with one worm per well. Each chemical condition (different colors) was tested across 2 columns of wells (n=12). Every plate contained one population of vehicle controls (white wells). Separate plates were tested for each planarian species. Worm drawing by Mackenzie Malia. B) Custom imaging setup allowing for recording of 2 plates simultaneously. Details of the components used are provided in Supplemental Table S2.

As shown in Figure 2B, recordings were taken with a custom imaging setup inside a NEEWER Photo Studio Shooting tent (Amazon.com, Seattle WA) that allows for recording in a controlled lighting environment. All components used in this set up are described in Supplemental Table S2. In brief, the plates were set on top of an LED light panel with brightness set to 1350 to 1450 lux, as measured with a lux meter. A Fresnel lens was placed on top of the plate to minimize imaging aberrations. While published studies primarily used 5-10 minutes of recording (e.g., (Bach et al., 2016; Pagán et al., 2015; Rawls et al., 2009) and up to 1 hour (Miller et al., 2025), we chose to record for 30 minutes, except where indicated, to allow for more sensitivity to detect pSLA that may not be induced immediately. Images were collected at 5 frames per second (fps) using a Point Grey Flea3 camera (FLIR, Wilsonville, OR) using Flycap software. This frame rate was chosen because we wanted to compare pSLA to other hyperactive behaviors, such as scrunching, which requires a minimum 5 fps frame rate for detection (Cochet-Escartin et al., 2015; Sabry et al., 2020), and we found that higher frame rates did not improve signal detection (Supplemental Figure S3).

### 2.4. Automated image analysis

Analysis of acute behavioral recordings was performed in MATLAB (2025a, MathWorks, Natick, MA) using custom scripts. Any worm that crawled out of the water during recording was excluded from analysis. To create a standardized image of the background to be using for masking the individual wells and removing background noise, the average projection of the first 2 minutes of recording was used. For determination of overall activity and changes in body shape, we calculated the absolute value of the difference of consecutive frames divided by the absolute value of the sum of the two images. The difference image was then binarized using a noise threshold determined from recordings of plates containing water only under the same imaging conditions. The motility index (MI) was then calculated for each well and frame as the sum of pixels that changed between the two images above the noise threshold. The area of each worm was calculated via thresholding of the image following subtraction by the average projection background image. As optical aberrations during imaging create some “blind spots” in which the planarian can become lost, any frames where no worm was found were excluded in the MI calculations. As individual bouts of pSLA have been previously described as lasting approximately 1 second (Rawls et al., 2009), the median MI score for 1 second (5 frame) bins was calculated. These scores were then normalized by the median worm area in the first 2 minutes of recording, where the worms are most likely to be moving the most and thus have elongated shapes.

To correlate normalized MI scores with the presence of pSLA, we manually evaluated the recording of 10-12 planarians exposed to NMDA (3 mM in GD, 15 mM in DJ) across 3 independent plates to identify a training set of manually determined pSLA events. For each analyzed worm, 1 minute of recording (60 1-second bins) was chosen, which exhibited high pSLA qualitatively by eye, and each 1 second bin containing pSLA was selected. For each species, we then evaluated the distribution of normalized MI scores across all identified pSLA events (Supplemental Figure S4). The 10^th^ percentile of each species’ pSLA distributions (GD: 0.11; DJ: 0.21) was then used as a threshold, where normalized MI scores above this threshold were counted as pSLA events.

MI provides information about the general activity of a planarian but gives no context to the spatial displacement of the worm. Thus, we also quantified the translational activity of each planarian over 20 seconds (100 frame) bins. This bin size was chosen because a normal gliding planarian can circle around the outside of a well of a 48-well plate in about this time. For each bin, the minimum intensity projection was determined. The average projection of the first 2 minutes of recording was subtracted from the minimum intensity projection to remove the background. The resulting image was binarized using a pre-determined threshold. The translational activity of each worm was calculated as the sum of pixels in this filtered image (thus representing the motion of the worm) divided by the median area of the worm during the first 2 minutes of recording. All image analysis codes have been deposited on Zenodo (doi: 10.5281/zenodo.18686049).

### 2.5. Lethality analysis

To determine if the presence of acute pSLA is correlated with subsequent lethality, some plates were removed after the initial acute recording and stored in a dark 20 °C incubator until the following day. Plates were recorded for 5 minutes at 21-27 hours after plate setup. From the recordings, each planarian was manually scored as alive, dead, “crawl-out” when the worm crawls out of the solution in the well, or “sick” when the worm was alive/moving but displayed abnormal morphology, indicated by head regression, pharynx extrusion, lesions/disintegration or C-shape body postures.

### 2.6. Statistical analysis

The number of pSLA events, defined by each species threshold (see section 2.4), over the entire 30 minute recording were counted and are provided in Supplemental File S1. Statistical analysis was performed in R (4.5.0, (R Core Team, 2016)). For each compound, data were compiled across all independent replicates and compared with the compiled in plate vehicle controls (planarian water or 0.5% DMSO) across the same replicates. A negative binomial generalized linear model was performed using the MASS package (Venables and Ripley, 2002) to determine an effect of condition on the number of pSLA events. Post hoc pairwise comparisons of the estimated marginal means of the model was performed using the contrast function of emmeans (Lenth, 2023) using the Benjamini & Hochberg (BH) p-value adjustment method for multiple comparisons. The median, 25^th^ and 75^th^ percentiles for each group, and final adjusted p-values for all pairwise comparisons are provided in Supplemental File S1.

## 3. Results

### 3.1. Defining pSLA events using prototypical concentrations of NMDA

The term pSLA was first introduced in 2009 by Rawls and co-authors to describe planarian behavior that looked convulsive and indicative of seizures in response to NMDA treatment (Rawls et al., 2009). However, pSLA identification was qualitative and the manual counting of pSLA is extremely time consuming and susceptible to bias. To overcome these limitations and thus make rapid screening of potential seizurogenic compounds in freshwater planarians possible, we developed a pipeline to robustly and quantitatively identify pSLA.

Because we needed a prototypical example of pSLA to build our analytical pipeline, we first identified a NMDA concentration that qualitatively induced robust pSLA in both planarian species within 30 minutes of exposure. Using the published data as a guide (Rawls et al., 2009), we tested 3 mM NMDA and found that this concentration immediately induced convulsive behavior in GD. However, robust convulsions were not observed in DJ until 15 mM NMDA was used. At these selected concentrations, NMDA-induced various paroxysms, including head flailing, turning, rapid twitching and convulsions, over-extension, C-shapes, corkscrews and spiral curling in both species (Supplemental Videos S1 and S2), as previously described in the literature. Figure 3 shows example high magnification images of the different shapes and behaviors that we observed in both species and Supplemental Figure S4 show the progression of behaviors over 15 minutes of recording for a *D. japonica* planarian exposed to NMDA.

**Figure 3.**
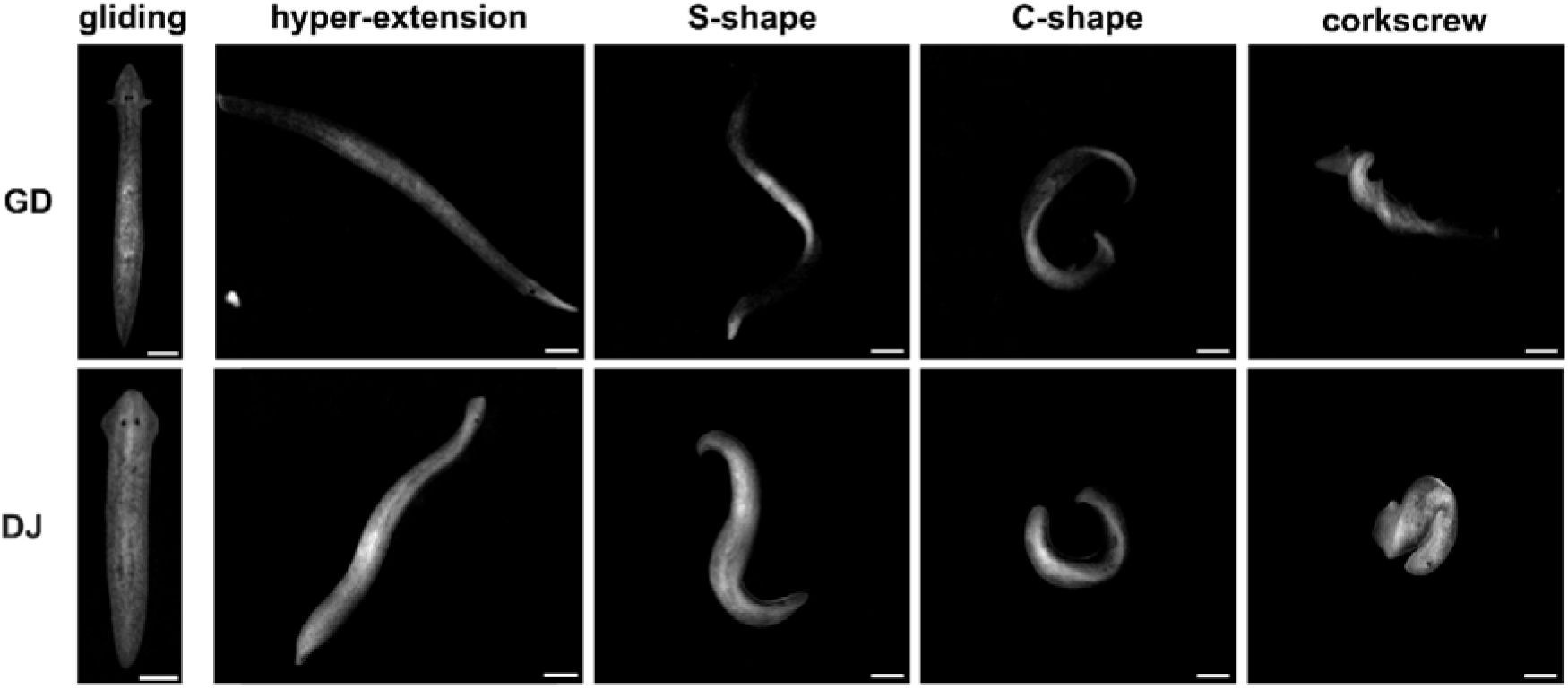
High magnification examples of abnormal body shapes and behaviors observed during NMDA-induced pSLA. Normal gliding behavior (left) as observed in planarian water exhibits relatively fixed body lengths and straight motion. During NMDA exposure (3 mM in GD and 15 mM in DJ), planarians change rapidly between different abnormal body shapes. Scale bars: 0.5 mm.

Next, to allow for rapid behavioral screening of multiple conditions simultaneously, we developed a medium-throughput screening set-up using 48-well plates as explained in Methods Section 2.3 and shown in Figure 2. Each well contained one planarian, with 2 columns per treatment condition, for a total of 12 planarians per condition per plate, allowing for simultaneous testing of 4 conditions in the same plate (Figure 2A). The use of commercial components (provided in Supplemental Table S2) for the behavioral imaging box (Figure 2B) allows for replication of this method in other laboratories. Using this set-up, chemically treated planarians were imaged for 30 minutes immediately (within 2-6 minutes) following exposure. To quantitatively identify pSLA, we measured the motility index (MI, see Methods section 2.4), normalized by worm area, for each planarian over the 30-minute recording period. As individual bouts of pSLA have been previously described as lasting approximately 1 second (Rawls et al., 2009), we calculated the median MI per 1 second (5 frames) bins for each worm. Species-specific pSLA thresholds were then determined based on the distribution of normalized MI scores for a subset of representative NMDA-exposed planarians (see Methods section 2.4, Supplemental Figure S5), where normalized MI scores per 1 second bin above this threshold were identified as potential pSLA events. We then applied this analysis to all NMDA data.

We found that NMDA showed significantly more pSLA events than the respective controls in each species (Figure 4A-B). Because these NMDA concentrations had low pH (∼3-3.6, Supplemental Table S1), we also tested the effect of pH alone using planarian water at approximately the same pH as the respective NMDA concentration. Acidic planarian water induced significantly more pSLA events than the respective non-adjusted controls, though not to the same extent as with NMDA. However, in DJ, the number of pSLA events in the acidic pH sample was not significantly different from NMDA. We conclude that pH is a contributing factor to NMDA-induced pSLA in both species.

**Figure 4.**
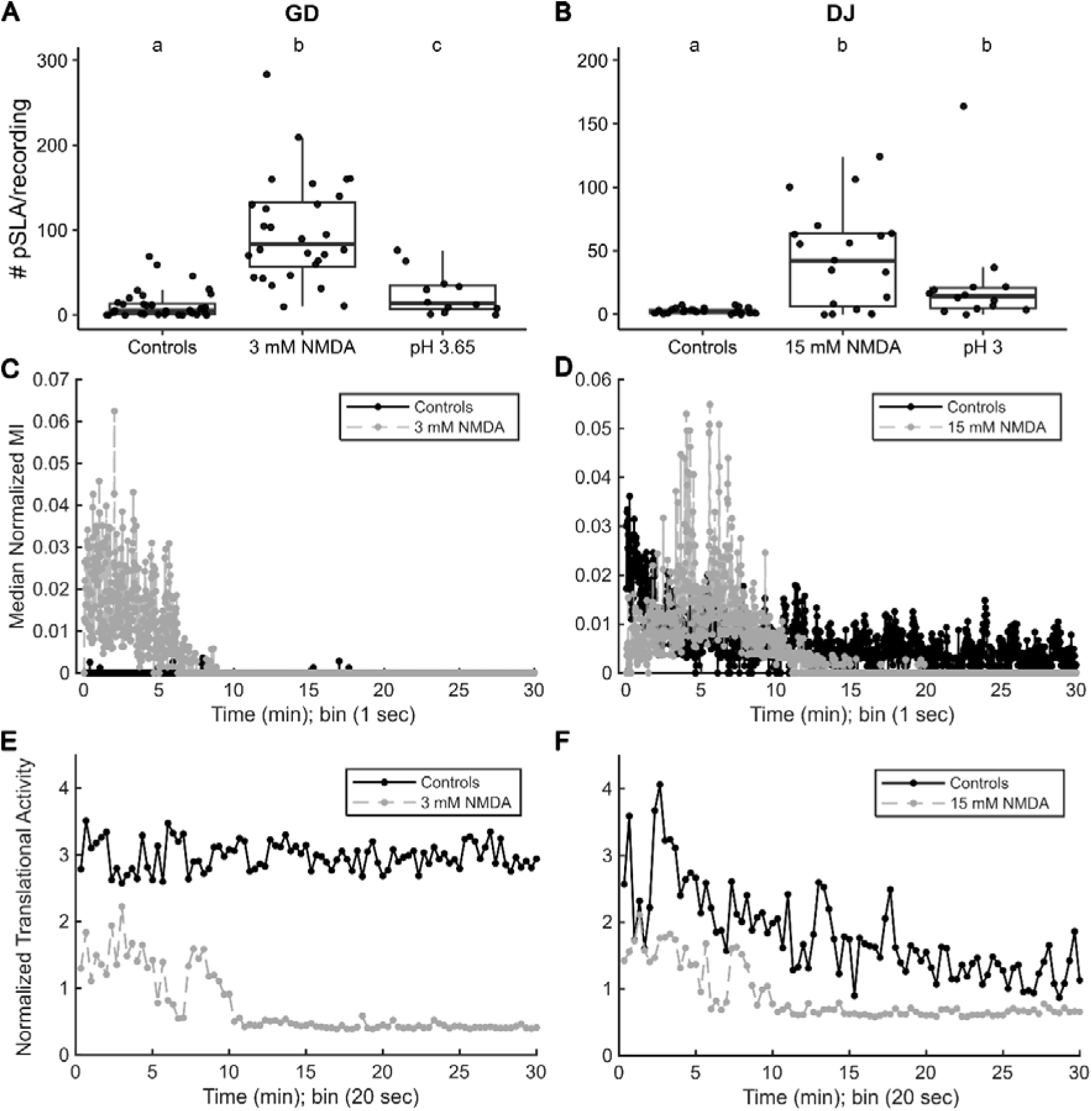
Validation of quantitative analysis of pSLA activity using NMDA. (A-B) Boxplots of the number of pSLA events per 30 min recording in A) GD and B) DJ exposed to NMDA and planarian water at the same pH as the respective NMDA concentration. Dots represent individual planarians. Statistical significance was determined using pairwise contrasts (with a Benjamini & Hochberg p-value correction) of the estimated marginal means of a negative binomial generalized linear model of condition versus number of pSLA events. Different lower-case letters indicate the groups are statistically significantly different (p<0.05). (C-D) Median normalized MI for all C) GD or D) DJ planarians exposed to their prototypical NMDA concentration and their respective in plate vehicle controls as a function of time. (E-F) Median normalized translational activity of all E) GD or F) DJ planarians exposed to their prototypical NMDA concentration and their respective in plate vehicle controls as a function of time. Sample sizes are listed in Supplemental Table S3.

While the total number of pSLA events over the entire recording time allows classification of a chemical as seizurogenic or not, there is additional information about the chemical’s action in the dynamics of the response, which can be evaluated from the normalized MI scores as a function of time (Figure 4C-D). In both species, NMDA induced high activity within 0-2 minutes of recording time, which dissipated within ∼5 (GD) or ∼10 (DJ) minutes. After this time, NMDA-exposed planarians became contracted and immobile and/or crawled out of the NMDA solution. As 0 or nearly 0 MI scores are seen with both normal gliding behavior and immobility, we also quantified the translational activity of the planarians (see Methods section 2.4). These translational activity plots show that the decrease in MI scores in NMDA-exposed planarians is concomitant with decreased motion around the well (Figure 4E-F). This decreased motility may indicate the toxic nature of NMDA as almost all worms in both species were dead or had crawled out of their wells by ∼24 hours (Supplemental Figure S6).

### 3.2. MI analysis distinguishes pSLA from other hyperactive behaviors

To verify that our analysis method is specific to pSLA, we exposed GD and DJ planarians to AITC, which has been shown to activate the planarian Transient Receptor Potential Ankyrin channel (DjTRPAa) and induce forward scrunching, a periodic muscle-based form of locomotion (Sabry et al., 2019). AITC concentrations that induced scrunching within a few minutes of exposure (25 µM for GD and 100 µM for DJ) had increased normalized MI scores (Figure 5A-B) and showed differences in translational activity compared to controls (Figure 5C-D). However, the MI scores were not high enough to be detected as significant pSLA activity (Figure 5E-F). AITC behavior was only analyzed for 10 minutes following exposure as scrunching did not persist past this time.

**Figure 5.**
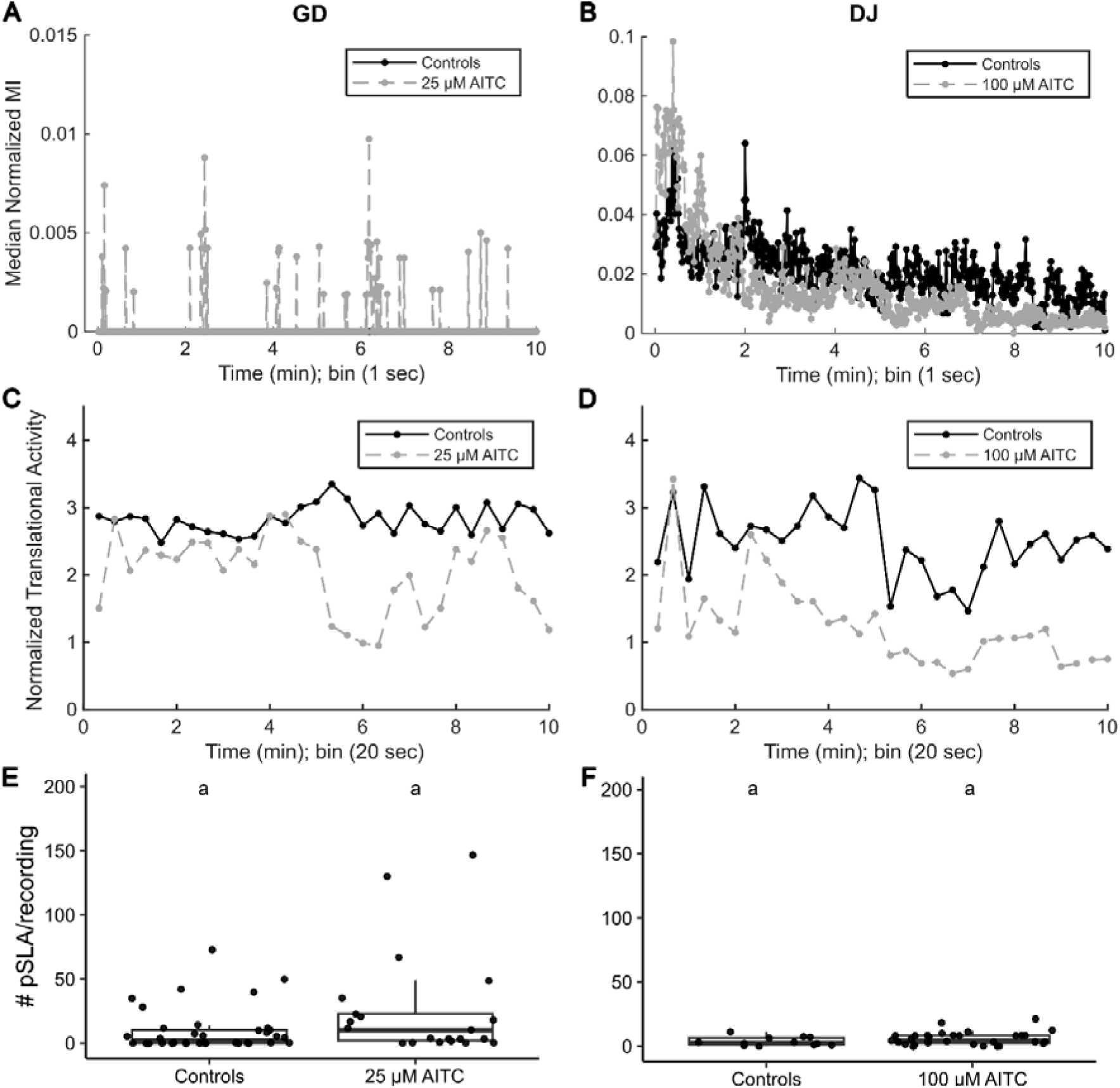
Quantitative MI analysis can distinguish pSLA from scrunching behavior induced by AITC. (A-B) Median normalized MI for all A) GD or B) DJ planarians exposed to AITC and their respective in plate vehicle controls as a function of time. Note that in A the control curve is not visible as all scores were 0. (C-D) Median normalized translational activity of all C) GD or D) DJ planarians exposed to AITC and their respective in plate vehicle controls as a function of time. (E-F) Boxplots of the number of pSLA events per 10 min recording in E) GD and F) DJ exposed to AITC and the respective in plate vehicle control. Dots represent individual planarians. No significant differences were found between controls and AITC in each using pairwise contrasts (with a Benjamini & Hochberg p-value correction) of the estimated marginal means of a negative binomial generalized linear model of condition versus number of pSLA events. Note the y-axes are scaled differently in the various plots to best reflect each data set. Sample sizes are listed in Supplemental Table S3.

### 3.3. Compounds known to induce pSLA in planarians are detected via MI analysis

Next, we tested other compounds that have previously been reported to induce pSLA in planarians (Table 2).

#### 3.3.1 Nicotine

Studies by different research groups have reported hyperactivity and pSLA in GD and GT planarians upon acute nicotine exposure in the µM to mM range (Figure 1 and Table 2). Nicotine binds to and activates nicotinic acetylcholine receptors. While there is supporting pharmacological evidence for nicotinic acetylcholine receptors in GD planarians (Rawls et al., 2011), the receptors have not yet been characterized. No significantly increased pSLA events were observed in GD planarians exposed to 10 – 50 µM nicotine (Figure 6A) although we qualitatively observed hyperactivity and pSLA-like events in GD planarians exposed to 10 µM nicotine immediately following exposure. This hyperactivity was slower and less intense than pSLA seen with 3 mM NMDA, as supported by lower amplitudes in the MI versus time plot for nicotine (Supplemental Figure S7) compared to NMDA (Figure 4C). In contrast to 3 mM NMDA, no lethality was observed at ∼24 hours with 10 µM nicotine (Supplemental Figure S6).

**Figure 6.**
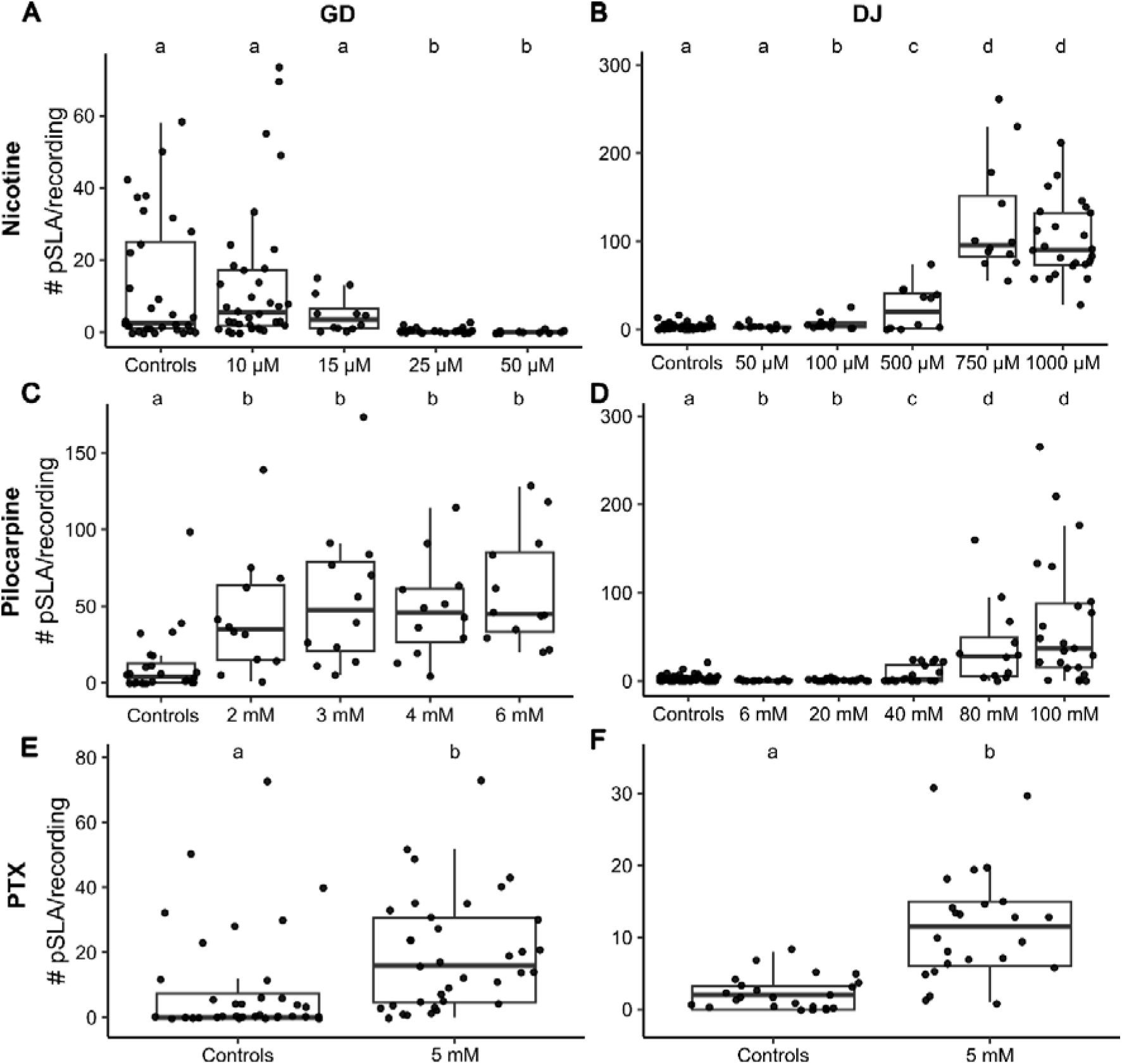
MI analysis of known seizurogenic compounds in planarians. Boxplots of the number of pSLA events per 30 min recording in (A-B) Nicotine, (C-D) Pilocarpine, and E-F) PTX in GD (left column, A, C, E) and DJ (right column, B, D, F). Dots represent individual planarians. Sample sizes are listed in Supplemental Table S3. Statistical significance was determined using pairwise contrasts (with a Benjamini & Hochberg p-value correction) of the estimated marginal means of a negative binomial generalized linear model of condition versus number of pSLA events. Different lower-case letters indicate the groups are statistically significantly different (p<0.05). Note the y-axes are scaled differently in the various plots to best reflect each data set.

Nicotine-induced pSLA quickly dissipated within the first 2 minutes, followed by a return to gliding behavior (Supplemental Figure S7 and S8). For these reasons, and because of the noise in the controls, no statistical significance was found when considering the number of pSLA events over the entire 30 minutes recording. When we focused on only the first 5 minutes of recording, we found significant pSLA for 10 µM nicotine (Supplemental Figure S9). Thus, our analysis shows that nicotine induces pSLA, in agreement with published data, which only scored 5 minutes of exposure (Ramakrishnan and DeSaer, 2011; Rawls et al., 2011).

Interestingly, pSLA was not observed in higher concentrations (15 and 25 µM) which largely displayed normal gliding. Exposure to 50 µM nicotine resulted in contraction and paralysis over the course of the 30 minutes recording (Supplemental Figure S10A). In contrast, in DJ planarians, significantly increased pSLA was observed starting at 100 µM nicotine, with the most pSLA events observed in 750 µM, which was highly hyperactive and demonstrated NMDA-like pSLA consisting of head flailing, convulsions and dynamic changes between S-and C-shapes (Figure 6B). In 750 µM nicotine, pSLA activity dissipated/slowed after 5 minutes of recording and was replaced by contraction with some sporadic, slow convulsive events (Supplemental Figure S11) and very little translational motion (Supplemental Figure S12). Contraction and further morphological abnormalities such as C-shapes and lesions were observed at ∼24 hours of exposure to 750 µM nicotine, although little lethality was observed (Supplemental Figure S6).

#### 3.3.2 Pilocarpine

It has recently been shown that a muscarinic receptor agonist, pilocarpine, can also trigger pSLA in GD. Using particle tracking to quantify the time spent in the center and the periphery of a well of a 12-well plate and manual scoring of different behaviors such as “dorsal oscillations”, “head flick”, “C-shape”, and others, concentrations of 4-6 mM pilocarpine were found to be significantly different from controls (Miller et al., 2025). Thus, we first tested 2-6 mM in GD planarians. Using MI analysis, we found that all tested concentrations of pilocarpine induced statistically significant pSLA in GD (Figure 6C), in agreement with the study by Miller et al. (Miller et al., 2025). No lethality was observed at 24 hours (Supplemental Figure S6). However, DJ planarians only displayed significantly increased pSLA at ≥ 40 mM pilocarpine, with the most pSLA events observed in 100 mM pilocarpine (Figure 6D), which also caused 100% lethality at 24 hours (Supplemental Figure S6). Thus, the two species show clear differences in sensitivity to pilocarpine, which may be due to differences in uptake, molecular targets, or metabolism.

#### 3.3.3 PTX

The GABA_A_ receptor antagonist PTX has been shown to induce pSLA in GT planarians (Ramakrishnan and DeSaer, 2011) but no studies have yet been conducted in GD or DJ planarians. Because of limited water solubility, the highest concentration of PTX that could be tested was 5 mM. We found that 5 mM PTX induced hyperactivity and pSLA within 30 minutes of recording in both species (Figure 6E-F). However, in contrast to the other chemicals tested, which induced pSLA within the first few minutes, 5 mM PTX induced pSLA was only observed after 20-25 minutes of recording, following initial gliding behavior and sporadic bouts of head twitching and dorsal oscillations (Supplemental Figures S7 and S11), suggesting that this concentration of PTX was at the potency threshold. PTX did not induce lethality in either species at ∼24 hours (Supplemental Figure S6), but the worms were elongated and were still exhibiting hyperactive behaviors.

### 3.4. PTZ can induce pSLA

Similar to PTX, PTZ also antagonizes the GABA_A_ receptor (albeit through a different mechanism) and is frequently used to generate animal seizure models (Afrikanova et al., 2013; Chitolina et al., 2023; Dhir, 2012; Milder et al., 2022; Shimada and Yamagata, 2018; Yang et al., 2023). We therefore tested varying concentrations of PTZ for its ability to induce pSLA in both GD and DJ planarians. We found that concentrations exceeding 10 mM (GD) or 5 mM (DJ) PTZ significantly induced pSLA (Figure 7A-B). PTZ-induced pSLA was characterized by an initial phase of head wiggling and turning, followed by random twitching and convulsions that included longitudinal over-extension and C-shapes, corkscrews and spiral curling. In GD, these hyperactive behaviors slowed down over the 30 minutes of recording and some of the planarians moved into a contracted state, indicating muscle paralysis (Supplemental Figures S7-8). In contrast, in DJ planarians, hyperactive behavior persisted throughout the entire 30-minute recording (Supplemental Figure S11). Both species were impacted at 24 hours with high numbers of death, sickness and crawl-out behavior, though the exact mix of these three abnormal conditions differed across the species (Supplemental Figure S6)

**Figure 7.**
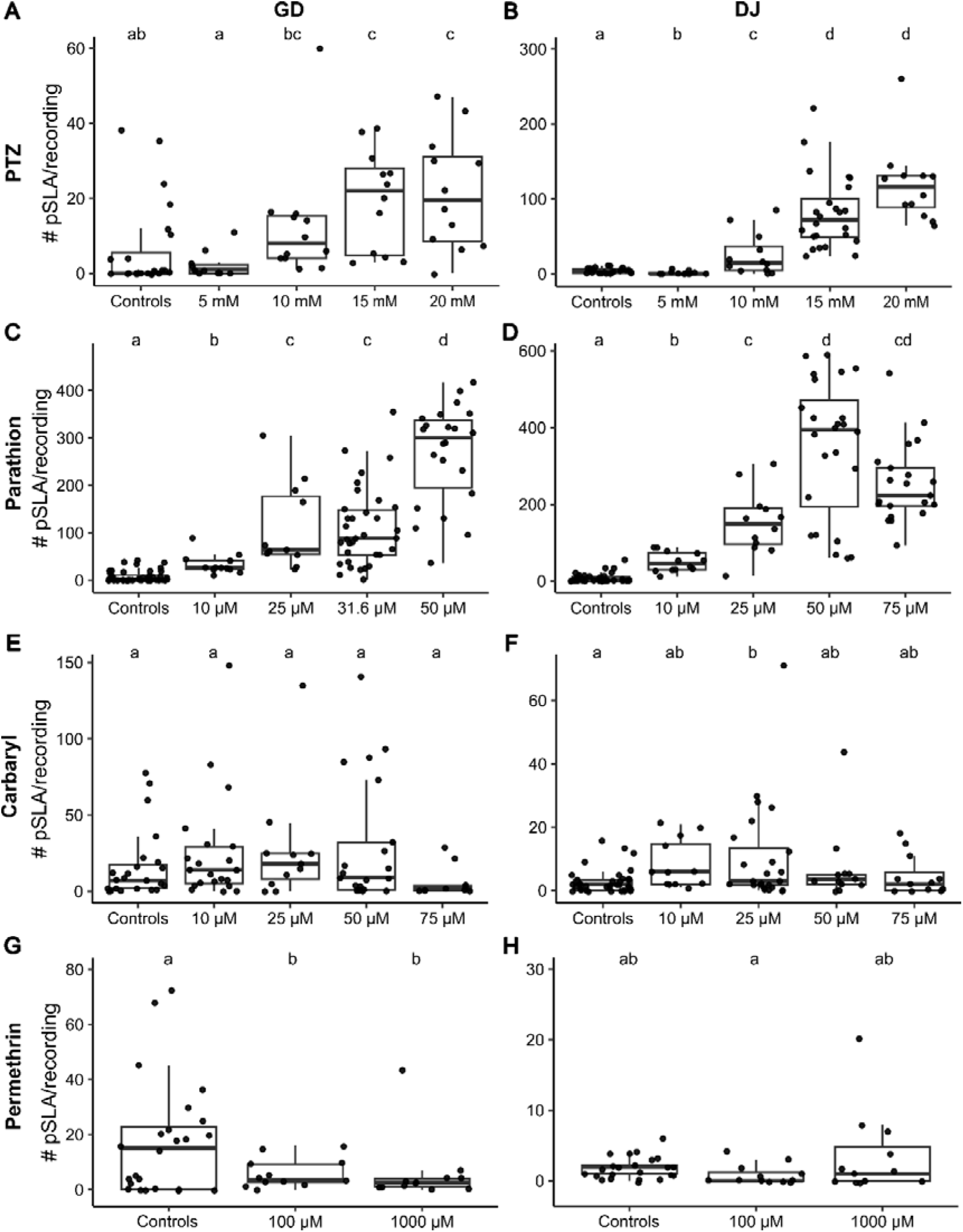
PTZ and parathion induce pSLA in GD and DJ planarians. Boxplots of the number of pSLA events per 30-minute recording in (A-B) PTZ, (C-D) parathion, (E-F) carbaryl, and (G-H) permethrin in GD (left column) and DJ (right column). Dots represent individual planarians. Sample sizes are listed in Supplemental Table S3. Statistical significance was determined using pairwise contrasts (with a Benjamini & Hochberg p-value correction) of the estimated marginal means of a negative binomial generalized linear model of condition versus number of pSLA events. Different lower-case letters indicate the groups are statistically significantly different (p<0.05). Note the y-axes are scaled differently in the various plots to best reflect each data set.

### 3.5. Some pesticides induce pSLA

Pesticides with different molecular targets have been found to be seizurogenic in mammalian models (Jett, 2012). Because both nicotine and pilocarpine, two cholinergic receptor agonists, induced pSLA, we hypothesized that organophosphorus pesticides and carbamates that inhibit acetylcholinesterase (AChE) in planarians as in humans (Hagstrom et al., 2017) and can cause seizures in mammals at high concentrations (Jett, 2012), could trigger pSLA upon acute high concentration exposure. To test this hypothesis, we tested one chemical from each class, parathion and carbaryl, respectively. Concentrations of 10 µM parathion and above caused significant pSLA in both planarian species (Figure 7C-D). While 50 µM caused the most pSLA events in both species, 31.6 µM parathion was a better concentration to induce pSLA in GD due to the high number of crawl-out events seen in this species at 50 µM. Following ∼24 hour exposure to these concentrations of parathion, both species showed sickness and lethality (Supplemental Figure S6). Carbaryl induced much weaker pSLA events in both species, which were characterized by small contractions and head twitches. While 25 µM showed significantly increased pSLA in DJ planarians compared to vehicle controls, 10-75 µM carbaryl showed no significant effect on the number of pSLA in GD planarians (Figure 7E-F), despite 25 µM showing increased median MI scores per bin (Supplemental Figure S7). At 75 µM, some scrunching and head twitching was observed within the first 5 minutes of recording, following by decreased movement and contraction (Supplemental Figure S10B), suggesting that this concentration exceeded the pSLA-inducing range of concentrations. No significant lethality was observed in either species exposed to 25 µM carbaryl at ∼24 hours (Supplemental Figure S6).

Some pyrethroid insecticides, such as permethrin, have been shown to cause seizures in rodents at lethal concentrations (Staatz and Hosko, 1985) or in environmentally relevant concentrations in cats (Dymond and Swift, 2008). Pyrethoids inhibit voltage-gated sodium channels, leading to neuronal hyper-excitability and at high doses can inhibit GABA signalling (Jett, 2012). We tested permethrin up to its solubility limit (1 mM) and did not observe significant pSLA in either planarian species (Figure 7G-H). Additionally, no obvious differences in behavior could be seen in the MI or translational activity plots (Supplemental Figures S7, S8, S11, S12).

### 3.6. The planarian brain is not required to induce pSLA in DJ planarians

It had previously been shown that pSLA in headless GT planarians was similar to that in intact planarians when exposed to nicotine but not to cocaine, another pSLA inducer (Pagán et al., 2012). Cocaine-induced pSLA was nearly completely lost in regenerating tails and was gradually restored during regeneration (Pagán et al., 2013). These results suggest that cocaine sensitive receptors are primarily found in the brain, consistent with dopamine receptors in vertebrates. In contrast, nicotinic acetylcholine receptors must be distributed in both the head and the body because the response in headless planarians was similar to that of intact planarians. Therefore, comparing responses of intact and headless planarians to seizurogenic chemicals may provide insights into where the molecular targets of these chemicals are expressed in planarians. Thus, we tested whether the presence of the brain was required for pSLA by exposing regenerating DJ tails to each seizurogenic compound at a concentration shown to significantly induce pSLA in adult/intact DJs. Exposure began 4 h after amputation to allow for wound closure and to mitigate the increased mucus production following amputation (Peiris et al., 2014) that may hinder chemical uptake. All chemicals except for carbaryl induced pSLA in regenerating DJ tails (Table 3).

**Table 3.** Median number of pSLA in adult or regenerating tail DJ planarians exposed to representative concentrations of seizurogenic compounds. Sample sizes are listed in Supplemental Table S3.

| Condition | Adult Median # pSLA (25th, 75th percentile) | Regenerating Median # pSLA (25th, 75th percentile) |
| --- | --- | --- |
| 15 mM NMDA | 42 (6, 63.5) | 45 (33, 68) |
| pH 3 | 14 (5, 21) | 69 (24, 132)* |
| 750 μM nicotine | 96 (83, 152) | 29 (11, 55)* |
| 100 mM pilocarpine | 37 (15, 88) | 69 (43, 201) |
| 5 mM PTX | 12 (6, 15) | 15 (10, 21) |
| 15 mM PTZ | 75 (49, 100) | 150 (93, 197) |
| 50 μM parathion | 394 (194, 470) | 59 (37, 136)* |
| 25 μM carbaryl | 3 (2, 13) | 0 (0, 1) |

\* non-overlapping interquartile ranges between adult and regenerating planarians for the same chemical

Some compounds, including NMDA, pilocarpine, PTX, and PTZ induced similar amounts of pSLA in the two developmental stages with overlapping interquartile ranges. Nicotine and parathion showed significantly more active behaviors - evidenced by higher numbers of pSLA - in adult DJ planarians than in tails, suggesting that the presence of the brain plays a role for pSLA induction in DJ planarians. Interestingly, pH 3 showed significantly greater activity in the regenerating tails than in adults. Thus, in DJ planarians, while the seizurogenic compounds do not strictly require the planarian brain to induce pSLA, presence of the brain can affect pSLA dynamics depending on the compound. Carbaryl was at the detection threshold and thus a comparison may not be meaningful.

## 4. Discussion

### 4.1. MI analysis can classify seizurogenic compounds and distinguish pSLA from other behaviors

To overcome the limitations associated with manual scoring of pSLA, namely scoring bias and time effort, we created an automated MI analysis method that uses image subtraction to detect changes in body shape. Using frame subtraction to quantify activity is a commonly used method to quantify behaviors including seizures in other model systems such as zebrafish (Afrikanova et al., 2013; Bruni et al., 2016; Whyte-Fagundes et al., 2025). By using qualitative scoring of NMDA-induced pSLA as a framework, we derived a quantitative MI threshold for pSLA in GD and DJ planarians. This analysis method was able to correctly identify known seizurogenic chemicals in planarians (NMDA, nicotine, PTX, pilocarpine) and distinguish pSLA from other abnormal behaviors, such as forward scrunching, which – in contrast to pSLA – exhibits periodic changes in body shape (Cochet-Escartin et al., 2015). Complementary analysis of the translational worm activity as a function of time helped us to distinguish normal behavior from contraction and paralysis, which both show low MI scores.

We confirmed previous studies by other research groups that NMDA (Rawls et al., 2009), nicotine (Rawls et al., 2011), and pilocarpine (Miller et al., 2025) induce pSLA in GD planarians and showed that they also induce pSLA in DJ planarians. PTX had previously only been studied in GT planarians (Ramakrishnan and DeSaer, 2011); here, we show that it also induces pSLA in GD and DJ planarians. The concentrations of NMDA, pilocarpine, and PTX that induced pSLA in GD planarians here were comparable to previous studies using GD or GT planarians. However, we found that 3 mM NMDA lowered pH to a range where pH alone caused pSLA while previous work found that 3 mM NMDA did not lower the pH as much (pH 6.6), and thus pH alone did not induce pSLA (Rawls et al., 2009). The discrepancy in these data may be due to differences in the exact water composition as the previous study used Artificial Pond Water (APW; 6 mM NaCl; 0.1 mM NaHCO_3_; 0.6 mM CaCl_2_; pH 7.3) (Rawls et al., 2009) which has slightly different salt composition compared to our planarian water (see Methods Section 2.1) and thus may have different buffering capacity.

For nicotine, we found that 50 µM caused paralysis in GD planarians, comparable to concentrations used in GT planarians (Ramakrishnan and DeSaer, 2011). In contrast, other studies in GD planarians have used nicotine in the mM range (Bach et al., 2016; Pagán et al., 2015, 2013; Rawls et al., 2011). The difference is likely be due to the product used. While we and (Ramakrishnan and DeSaer, 2011) used pure (-) nicotine solution, the other studies (Bach et al., 2016; Pagán et al., 2015, 2013; Rawls et al., 2011) used nicotine hydrogen tartrate salt. The pesticides parathion and carbaryl are both cholinesterase inhibitors and also induced pSLA in at least one planarian species whereas the pyrethroid permethrin, which inhibits voltage-gated sodium channels, did not.

Finally, we showed that PTZ, which is used to generate seizure models in mice (Dhir, 2012; Shimada and Yamagata, 2018) and zebrafish (Afrikanova et al., 2013; Baraban et al., 2005; Chitolina et al., 2023; Milder et al., 2022), robustly induces pSLA in both planarian species. Thus, PTZ could be used to generate a planarian seizure-like model to test the efficacy of possible anti-seizure treatments. In fact, existing studies that used compounds other than PTZ suggest that pSLA in planarians can be modulated by exposure to known anti-convulsant therapeutics (e.g., (Pagán et al., 2015; Ramakrishnan and DeSaer, 2011; Rawls et al., 2009)). Now that we have developed this automated platform, it can be used to validate existing manual studies on therapeutic effectiveness.

### 4.2 GD and DJ planarians respond similarly to chemicals but with different sensitivities

GD and DJ planarians displayed very different sensitivities to NMDA, nicotine, and pilocarpine (Table 3, Supplemental Tables S4-S5). GD planarians were more sensitive, with the most extreme difference in sensitivity of nearly 100-fold for nicotine. While NMDA and pilocarpine induced similar behavioral phenotypes in the two species, there were also qualitative differences in the behaviors displayed by the two species in response to nicotine as DJ planarians exhibited extensive pSLA events that were never observed in GD planarians. Other studies have reported similar behavioral and sensitivity differences between planarian species in response to certain chemicals (e.g., (Ireland et al., 2020; Sabry et al., 2019)). Together, these data emphasize that extrapolation between planarian species is not recommended due to possible differences in pharmacokinetics (chemical uptake, metabolism) and/or pharmacodynamics (presence of targets/receptors). Further mechanistic studies will be needed to better understand these species-specific differences.

We found that DJ planarians are more behaviorally heterogeneous than DJ planarians. These differences may be due to the DJ planarians being bred under controlled laboratory conditions with standardized food and culture conditions for years whereas GD planarians had only been maintained under these conditions for a few weeks in our study. Changes and differences in diet could also contribute to the behavioral differences (Pacis et al., 2025). The variability in GD behavior was the primary reason for reduced statistical power in some of the exposure conditions (Supplemental Table S6). However, the increased sensitivity of GD planarians to some of the compounds tested is a strength for screening seizurogenic chemicals. More compounds will need to be screened in both species to evaluate their strengths and weaknesses for this application. A good starting point would be to test chemicals that have been proposed to induce pSLA in planarians in recent manual studies, such as sertraline (Thumé and Frizzo, 2017) and various cholinesterase inhibitors (Bezerra Da Silva et al., 2016), which were not included here.

### 4.3 The brain is not required for pSLA

Most seizurogenic chemicals induced pSLA in headless DJ planarians to a similar extent as in intact adults, indicating that the cephalic ganglion is not required for pSLA induction by the compounds tested here. The planarian CNS is distributed and includes both the cephalic ganglion and two ventral nerve cords (VNCs) that extend along the head–tail axis (Ross et al., 2017). Thus, hyperexcitation of neural circuits within the VNCs may be sufficient to generate pSLA. This parallels observations in mammals that spinal cord circuits can independently support seizure-like activity. For example, GABA_A_ receptor blockade with PTX induces epileptiform activity in mouse spinal neurons *in vitro* (Barker and MacDonald, 1980). These findings suggest that neural circuits can generate hyperexcitability even without brain involvement.

Given that planarians reproduce asexually through binary fission, autonomous motor control of a headless body fragment is essential for survival of tail offspring (Le et al., 2021). The persistence of pSLA following decapitation may therefore reflect functional redundancy within this distributed neural architecture rather than the absence of a neural origin. Consistent with this view, planarian behavioral responses are complex and many motor behaviors — including turning, rectifying, and scrunching — can occur in the absence of the head (Cochet-Escartin et al., 2015; Le et al., 2021), indicating that the VNCs can independently coordinate motor output. However, our data do not allow us to distinguish whether the observed effects arise from central versus peripheral nervous system mechanisms, nor whether they reflect neuronal hyperactivity or direct activation of muscle. Future mechanistic studies (see 4.5.3) will be needed to distinguish these scenarios.

Interestingly, parathion and nicotine both produced significantly more pSLA events in intact adult *D. japonica* than in headless planarians, suggesting that the brain enhances or modulates the behavioral response to these compounds. Although cholinesterases (Hagstrom et al., 2018) and nicotinic receptors (Pagán et al., 2013) are distributed throughout the planarian body, cholinesterase expression is highest in the brain, indicating that cholinergic signalling may be more strongly regulated there. The cephalic ganglion may therefore amplify or coordinate cholinergic network activity, leading to more robust pSLA when intact. Consistent with a modulatory role for the brain, nicotine exposure previously produced fewer pSLA events in decapitated GT planarians, although the effect was not statistically significant (Pagán et al., 2013). Differences between that study and ours may reflect species differences, nicotine formulation, or analytical methods. Notably, cocaine-induced pSLA has also been reported to require the presence of the brain (Pagán et al., 2013), further supporting the idea that at least some substances depend on cephalic neural integration to produce maximal behavioral output. Because our analysis compared only the medians and interquartile ranges of the total number of pSLA events over the 30-minute recording, differences in response dynamics between intact and headless planarians would not have been detected and may warrant future investigation.

### 4.4 Use of planarians for screening seizurogenic chemicals and therapeutics

Planarians offer practical advantages for early seizure liability screening. Although they are simpler than zebrafish and lack a circulatory system and blood–brain barrier (Rawls et al., 2010), they are inexpensive to maintain and, as invertebrates, do not require the animal welfare protocols mandated for vertebrate models like zebrafish (Collins et al., 2024). Zebrafish larvae at 5 days post fertilization (dpf) are primarily used for screening when experiments are not classified as animal testing (Strähle et al., 2012) but their nervous system and metabolic machinery are still maturing (Menezes et al., 2014; Saad et al., 2017; Tiedeken and Ramsdell, 2010). Seizure models have also been developed for *Caenorhabditis elegans*, but rely on specific genetic strains and have largely been used for identification of anti-seizure therapeutics (Jones et al., 2021; Wong et al., 2018). We envision a drug discovery pipeline where, following *in silico* chemical modelling and/or human *in vitro* MEA assays, compounds can be screened in adult planarians with metabolic capacity (Ireland et al., 2022a) and mature neuronal function, before subsequent testing in developing zebrafish and preclinical rodent models. This tiered screening approach would only prioritize the best lead candidates at each stage, ultimately reducing cost and vertebrate use in early seizure liability testing. We have previously shown that organismal planarian and zebrafish behavioral screening and MEA assays were the most informative to identify developmental neurotoxicants (Ireland et al., 2025b) and thus may prove to be a promising combination for identify seizurogenic chemicals and anti-seizure therapeutics.

All models have limitations, some of which are unique and some which are shared across models. The strengths and limitations of planarians and developing zebrafish (<5 dpf) for chemical screening have recently been reviewed (Collins et al., 2024). One limitation shared by organismal models and cell culture models is the requirement for aqueous chemical exposure, making poorly water-soluble or volatile compounds difficult to test (Collins et al., 2024). As a result, sufficiently high concentrations to induce pSLA may not always be achievable, as illustrated by our ability to test PTX only at its seizure induction threshold. *In silico* modelling can help overcome some experimental limitations, as has been shown recently for screening anti-seizure drugs in zebrafish larvae (Belair et al., 2025; Cassar et al., 2017). Comparative testing of compounds known to be seizurogenic in humans across these models can clarify their complementarity within a rapid seizure liability test battery and identify data gaps and limitations, similar to ongoing efforts in developmental neurotoxicity testing (Ireland et al., 2025b; Masjosthusmann et al., 2020).

### 4.5 Limitations and Opportunities

The work here can act as a starting point for future screens of seizurogenic chemicals and anti-convulsants in planarians but further optimization and system validation are required to decide whether planarian screening could be a valuable part of an early-stage seizure liability testing battery. Three major tasks still remain: 1) optimized experimentation to improve throughput, 2) optimized image analysis and 3) mechanistic validation of pSLA.

#### 4.5.1 Optimize experimental throughput

The experimental approach described here provides a proof-of-concept design that should be further optimized to fit within the needs of a preclinical drug discovery pipeline. The recording period could be shortened to 15 minutes to achieve higher throughput as most chemicals induced pSLA within the first 5-10 minutes. However, it should be recognized this may decrease detection power for some chemicals. Our recordings began 2-6 minutes following chemical exposure due to experimental constraints in starting recording immediately, which was sufficient for the compounds we studied (Supplemental Figure S13). This window could be decreased to increase the possibility of catching very short-lasting pSLA events. Here, we used a minimum of n=12 specimen per condition, similar to or greater than the sample sizes used in previous planarian seizure studies (Bach et al., 2016; Miller et al., 2025; Rawls et al., 2009). While these sample sizes were generally sufficient to achieve adequate statistical power, intrinsic variability in control behavior of GD planarians required larger sample sizes in some cases (Supplemental Table S6). Therefore, while the column setup that we used here, utilizing n=12 within a single experiment, was helpful for testing a few selected compounds for this proof-of-concept study, for new screens, we recommend a row setup, similar to what is used for rapid behavioral screening in other studies (Ireland et al., 2025a; Zhang et al., 2019), because it allows for the simultaneous testing of 5 concentrations/conditions plus a vehicle control. By using 2 plates with alternating row setup, one can control for potential edge effects (Zhang et al., 2019) and achieve n=16 per condition while testing in independent experiments to ensure robustness. In addition to acute behavioral recordings, we also evaluated lethality one day after chemical exposure to determine if lethality was correlated with the presence of significant pSLA. This was not the case. Thus, lethality likely results from other toxic effects besides pSLA and thus is not an informative endpoint for this type of study. To accelerate system evaluation and have multiple research groups participate in characterizing the value of planarian behavioral screening, the recording and analysis methods developed here are made available to other laboratories. This will ensure standardized conditions that allow cross-laboratory data comparisons and integration.

#### 4.5.2 Optimize automated image analysis

The MI analysis pipeline described here provides an automated, quantitative way to detect pSLA that improves on previous manual techniques that were not well defined and tedious. One limitation of MI analysis is that it has difficulty to detect pSLA when the convulsive/twitching behavior is small and/or primarily in the third dimension (e.g., in the case of carbaryl), which is not well captured in the simple 2D imaging setup we used. While these behaviors could still be detected as pSLA in DJ planarians, they were not found to be significant for GD planarians because GD planarians showed more variable control behavior, including swimming, scrunching, and turning over. Future work could incorporate more sophisticated image analysis methods, incorporating machine learning to identify specific body shape changes and use them to characterize pSLA, as has recently been done in zebrafish (Whyte-Fagundes et al., 2025). Such a tailored approach would likely be more robust than MI analysis and may reveal subtypes of pSLA, which could provide additional insight into the underlying biological mechanisms. For example, the initial behavioral effects of parathion are thought to be due to acetylcholinesterase inhibition while later effects are from secondary neurotransmitter systems such as glutamate (Jett, 2012; Kozhemyakin et al., 2010); thus, the type of convulsions may differ over time. Data analysis could thus also consider the temporal dynamics of the behavioral response by evaluating effects over different time bins.

#### 4.5.3 Mechanistic validation of pSLA as a model for human seizure activity

For planarians to be used as a screening system for seizure liability, it is crucial to confirm that pSLA is indicative of seizures using electrophysiological measurements of neuronal activity. Some electrophysiological techniques have been developed in planarians (Aoki et al., 2009; Benita et al., 2024; Freiberg et al., 2023), with varying levels of invasiveness. However, existing techniques are relatively low throughput and not well suited to chemical exposure. Alternatively, MEA has been successfully used to measure neuronal activity *in vivo* in zebrafish larvae (Tomasello and Sive, 2020) and could be adapted for planarians. In addition, RNA interference (RNAi) could be used to verify that the known seizurogenic compounds tested here induce pSLA through the same pathways as in mammals. For example, PTX and PTZ both target the GABA_A_ receptor, thus making it a good starting point for mechanistic studies. Both parathion and carbaryl induced pSLA in DJ planarians and inhibit DJ cholinesterases (Hagstrom et al., 2017; Ireland et al., 2022b); thus, RNAi knockdown of the planarian cholinesterases could be used to verify that the molecular initiation event of pSLA in planarians is cholinesterase inhibition, as in humans (Jett, 2012). These verification studies will be critical for adoption of the planarian model for screening seizurogenic chemicals as they would show that results found in planarians are relevant and translatable to mammals and humans.

In summary, this work demonstrates that pSLA can be triggered through various pathways in GD and DJ planarians, similar to in mammals. By standardizing both experimental approach and analysis methods and making the tools available to the research community, this work can serve as a framework for future studies of chemicals for their seizurogenic potential in planarians.

## CRediT authorship contribution statement

**Danielle Ireland:** Conceptualization; Software; Visualization; Validation; Formal analysis; Data curation; Writing – original draft; Writing - review and editing. **Evangeline Coffinas:** Investigation; Writing – review and editing. **Christina Rabeler:** Investigation; Writing – review and editing. **Eva-Maria S. Collins:** Conceptualization; Software; Methodology; Visualization; Validation; Writing – original draft; Writing – review and editing; Supervision; Resources; Project administration.

## Data availability

The datasets supporting the conclusions of this article are included within the article and available from the corresponding author upon reasonable request. The MATLAB code used for image analysis is available on Zenodo (doi: 10.5281/zenodo.18686049).

## Acknowledgements

The authors thank Kate Sun and Erin Szuromi for their help with experiments, Dr. Scott Rawls, Dr. Hilary McCarren, and Mathieu Conroy for discussions, Mackenzie Malia for the worm schematic in Figure 2A and Hannah Poon for comments on the manuscript. During the preparation of this work, the authors used ChatGPT (OpenAI) in order to help design R code for the power analysis presented in Supplemental Table S4. After using this tool, the authors reviewed and edited the content as needed and take full responsibility for the content of the published article.

