## Supplementary Figures and Tables for "Planarian behavioral screening is a useful invertebrate model for evaluating seizurogenic chemicals"

### Supplemental Figures

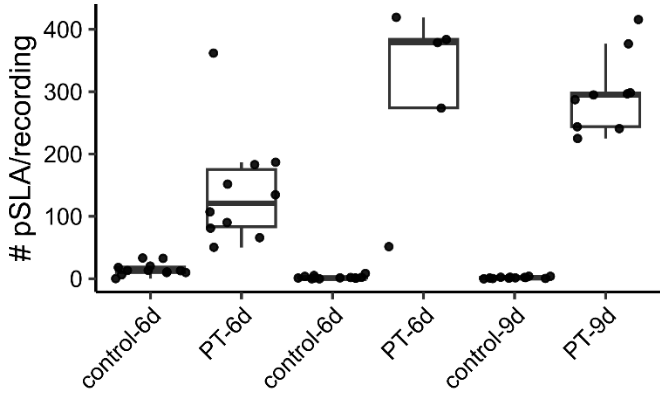


**Supplemental Figure S1. Length of planarian fasting does not drastically change results.** Boxplots show the number of pSLA events for DJ planarians exposed to 0.5% DMSO (control) or 50 µM parathion (PT). Data are shown for each tested plate separately and list the number of days (d) the planarians were fasted. Each pair of control and PT represent one plate. While there is plate to plate variability in the number of pSLA events, differing fasting durations do not dramatically change the results. Sample sizes are listed in Supplemental Table S3.

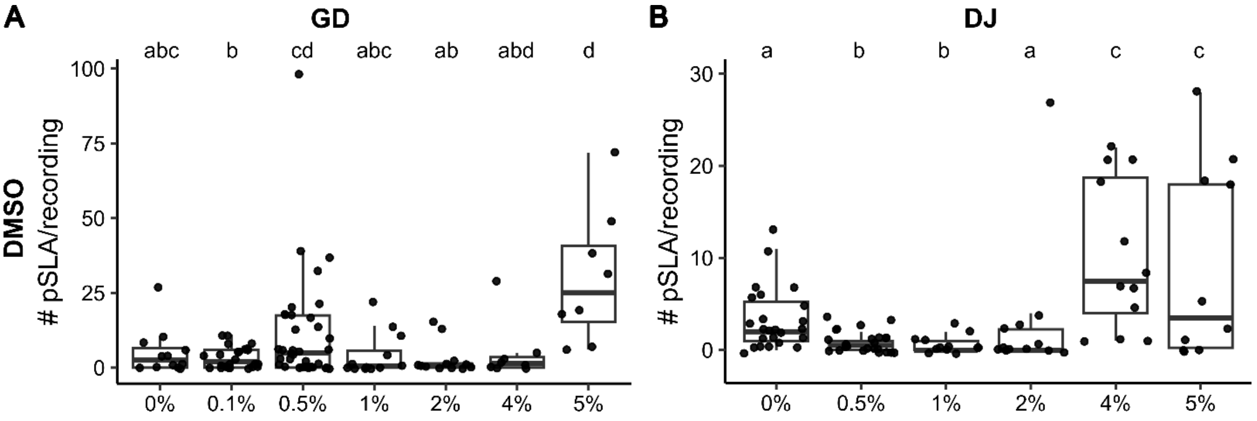


**Supplemental Figure S2. DMSO concentrations less than 4% do not cause pSLA.** A) GD or B) DJ planarians were exposed to varying concentrations of DMSO. The number of pSLA events were counted using the Motility Index pipeline as described in Methods Section 2.4. 0% refers to untreated planarian water. Significantly increased number of pSLA events were only observed in 5% DMSO in GD or 4 and 5% in DJ. Statistical significance was determined using pairwise contrasts (with a Benjamini & Hochberg p-value correction) of the estimated marginal means of a negative binomial generalized linear model of condition versus number of pSLA events. Different lower-case letters indicate the groups are statistically significantly different ($p<0.05$). Sample sizes are listed in Supplemental Table S3.

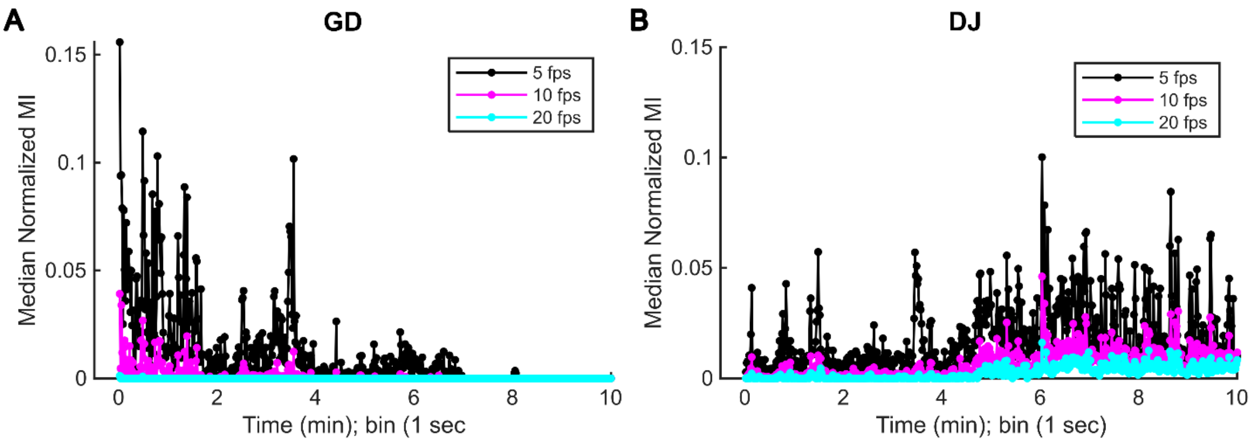


**Supplemental Figure S3. A frame rate of 5 fps is sufficient to detect pSLA.** One plate testing 3 mM NMDA in GD planarians (A) or 15 mM NMDA in DJ planarians (B) was recorded for 10 minutes at 20 frames per second (fps). MI analysis was performed using the entire dataset or down sampling of the original video to mimic 10 or 5 fps datasets. Data show the median response of n=12 (GD) or n=6 (DJ). Higher frame rates result in smaller MI scores and generally less resolution of pSLA.

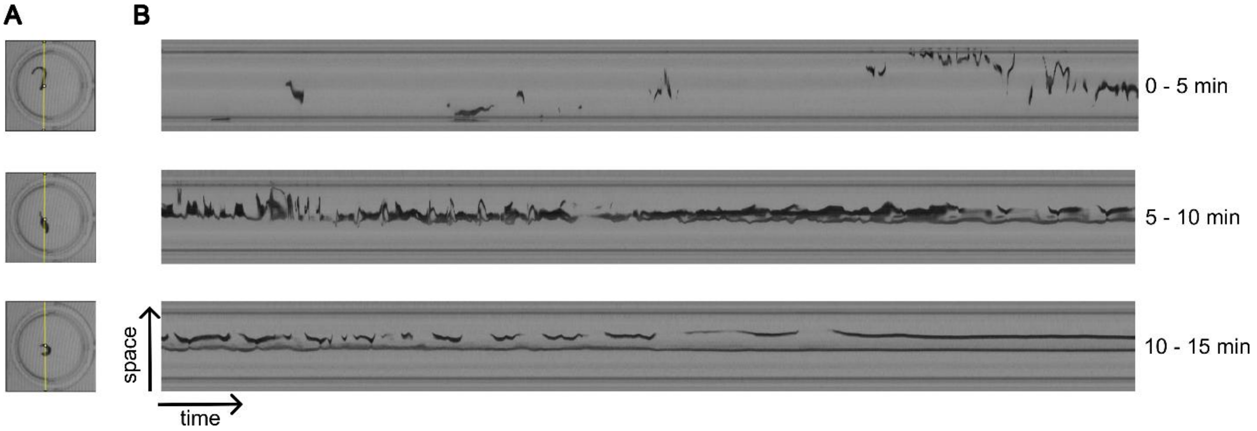


**Supplemental Figure S4. Representative kymograph to show pSLA progression.** Video of a DJ planarian exposed to 15 mM NMDA was used to show a representative progression of pSLA. A) Line (in yellow) indicates slice where kymograph was taken. B) Each kymograph represents motion in space (y-axis) over 5 minutes of activity (x-axis). The worm is initially gliding and then begins small shape changes which turn into pSLA around 4 minutes of recording. Eventually the shape changes become smaller in amplitude and occur less frequently until the worm stops moving (~14 minutes onward).

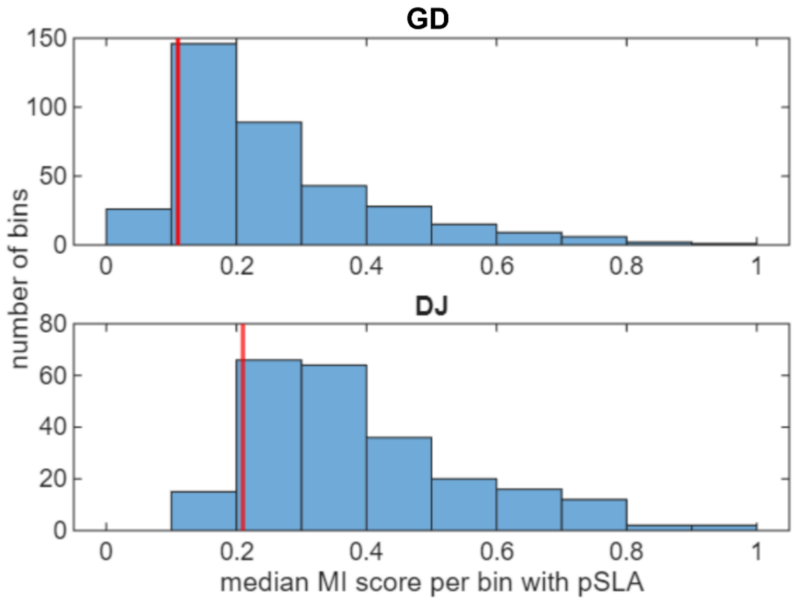


**Supplemental Figure S5. MI scores of NMDA-induced pSLA events.** Recordings of NMDA-induced behavior (3 mM in GD; 15 mM in DJ) were manually scored to identify 1 second bins containing pSLA events, defined as rapid body shape changes. Manual scoring was done on 10-12 worms per species across 3 independent experiments and for approximately 1 minute (60 bins) per worm. The scored minute was qualitatively chosen as a highly active minute with many pSLA events. The distribution of the median normalized MI score per 1 second bin for these pSLA events are shown for each species. For subsequent analysis, a pSLA event was identified as a 1 second bin with a median normalized MI score greater than the $10^{th}$ percentile of the NMDA distributions shown above (red lines, GD: 0.11, DJ: 0.21).

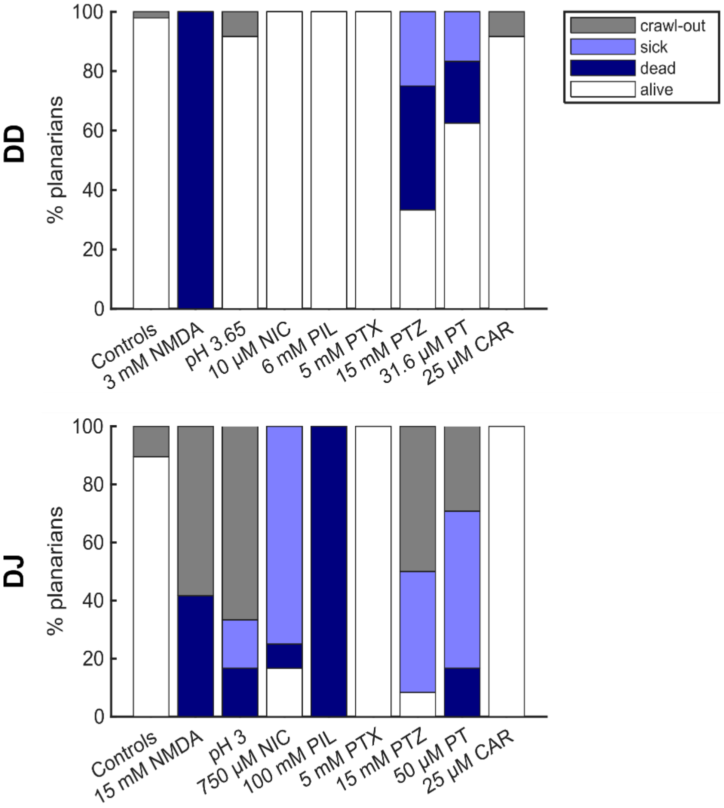


**Supplemental Figure S6. Lethality of pSLA inducing concentrations after ~24 hours of exposure.** Select plates of GD (top) and DJ (bottom) planarians exposed to pSLA-inducing concentrations of the seizurogenic compounds were re-evaluated at ~24 hours (21-27 hours) following continuous exposure. Note the difference in concentrations for GD and DJ planarians for some of the compounds. Lethality/morphology was manually scored from 5 minute recordings. Planarians were classified as sick if they were alive but had lesions, partial disintegration, pharynx extrusion, head regression or immobile C-shapes. Crawl-out refers to worms that crawled out of the well in the water. NIC: nicotine, PIL: pilocarpine, PT: parathion, CAR: carbaryl. Sample sizes are listed in Supplemental Table S3.

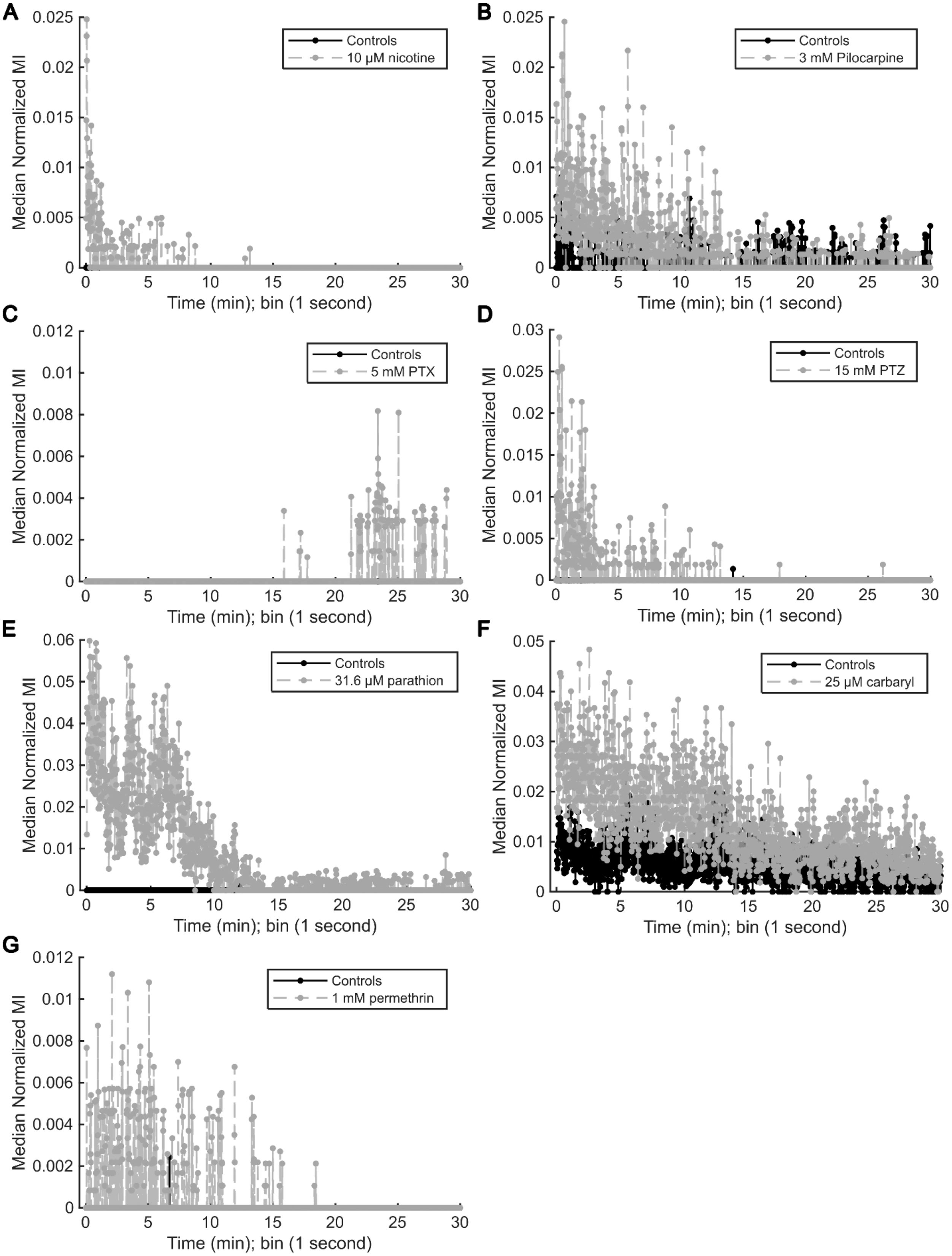
A
B
C
D
E
F
G
Median Normalized MI
Time (min); bin (1 second)
Controls
10 μM nicotine
3 mM Pilocarpine
5 mM PTX
15 mM PTZ
31.6 μM parathion
25 μM carbaryl
1 mM permethrin

**Supplemental Figure S7. MI versus time plots for select concentrations of chemicals in GD planarians.** The median MI of all tested planarians for each 1 second bin are shown for the concentrations with the highest number of median pSLA for each chemical compared to its in-plate vehicle control. Note the y-axes are scaled differently in the various plots to best reflect each data set. Some control activity that consists of only 0 median MIs are not visible. Sample sizes are listed in Supplemental Table S3.

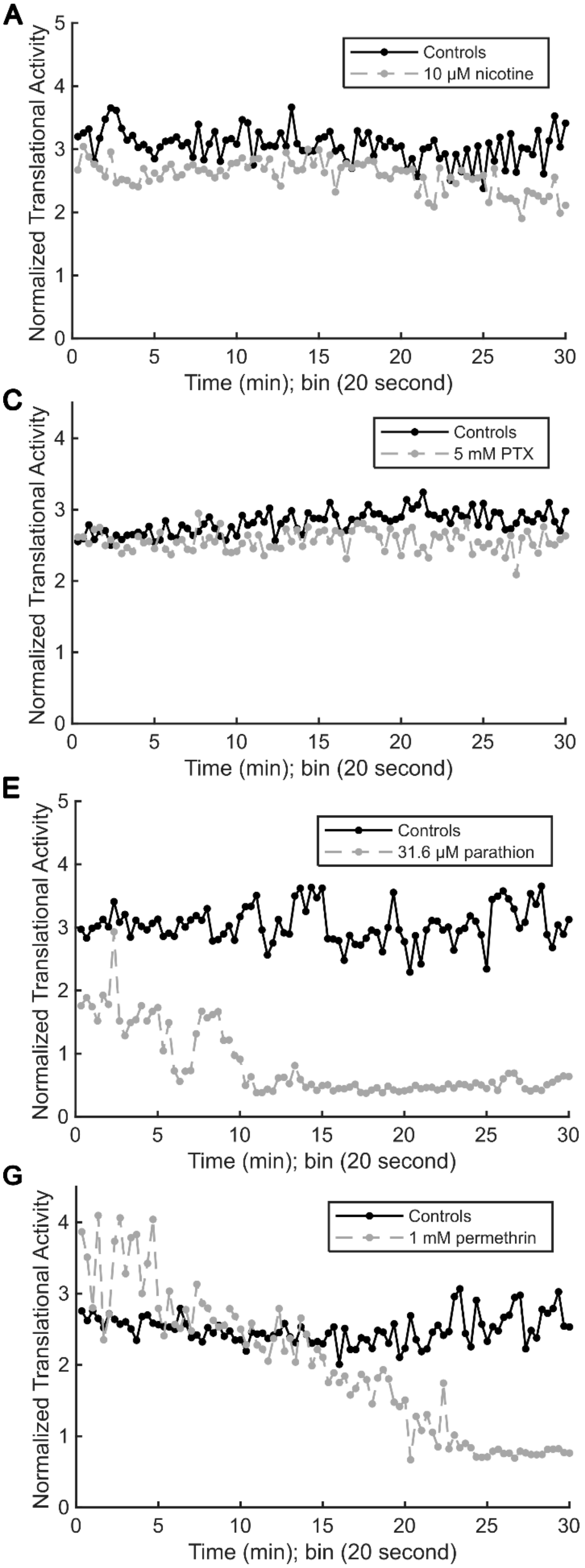
A
Controls
10 μM nicotine
B
Controls
3 mM pilocarpine
C
Controls
5 mM PTX
D
Controls
15 mM PTZ
E
Controls
31.6 μM parathion
F
Controls
25 μM carbaryl
G
Controls
1 mM permethrin
Normalized Translational Activity
Time (min); bin (20 second)

**Supplemental Figure S8. Translational activity vs time for select concentrations of chemicals in GD planarians.** The median normalized translational activity of all tested planarians for each 1 second bin are shown for the concentrations with the highest number of median pSLA for each chemical compared to its in-plate vehicle control. Note the y-axes are scaled differently in the various plots to best reflect each data set. Sample sizes are listed in Supplemental Table S3.

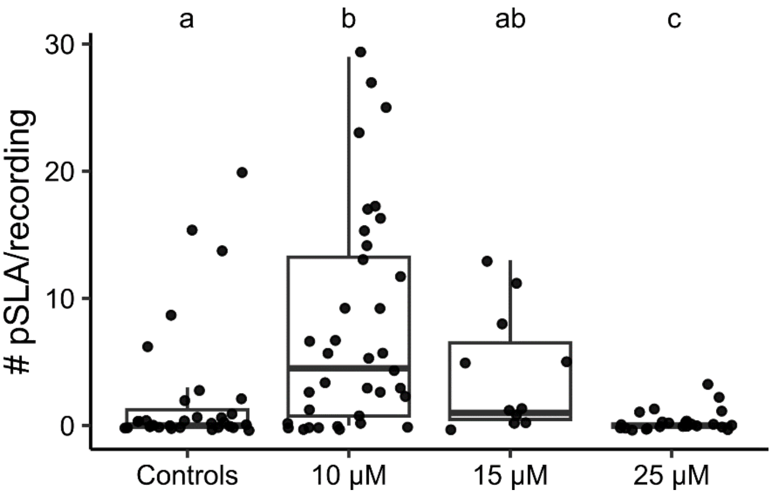


**Supplemental Figure S9. Nicotine causes significant pSLA in shorter time intervals.** Boxplots of the number of pSLA events in the first 5 minutes of recording in GD planarians shows significant pSLA in 10 μM nicotine. 50 μM is not shown as it consisted of only 0 pSLA events (n=12). Statistical significance was determined using pairwise contrasts (with a Benjamini & Hochberg p-value correction) of the estimated marginal means of a negative binomial generalized linear model of condition versus number of pSLA events. Different lower-case letters indicate the groups are statistically significantly different ($p<0.05$). Dots represent individual planarians. Sample sizes are listed in Supplemental Table S3.

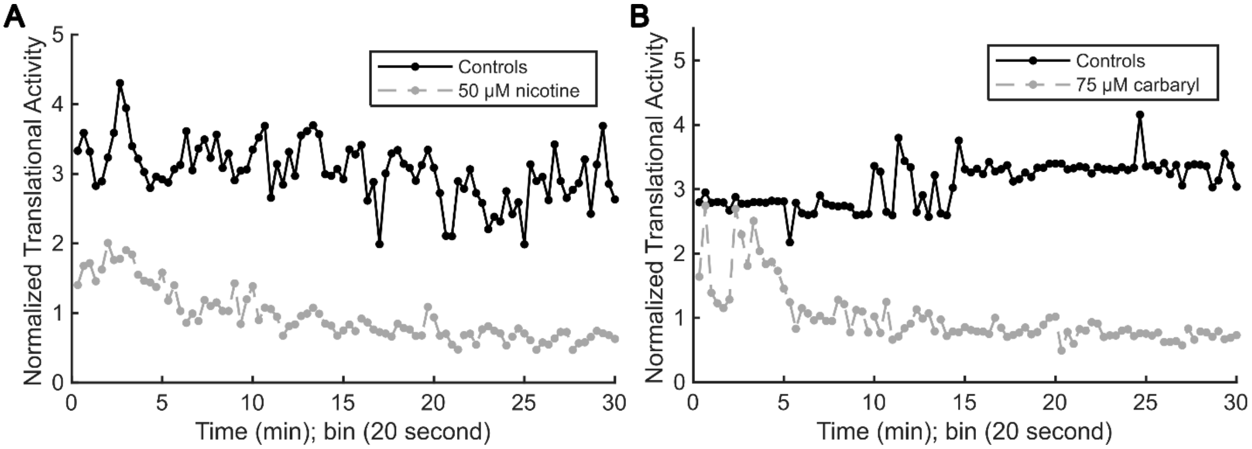


**Supplemental Figure S10. Translational activity vs time for conditions that induce contraction and immobility in GD planarians.** The median normalized translational activity of all tested planarians for each 1 second bin are shown for the concentrations with the highest number of median pSLA for each chemical compared to its in-plate vehicle control. Sample sizes are listed in Supplemental Table S3.

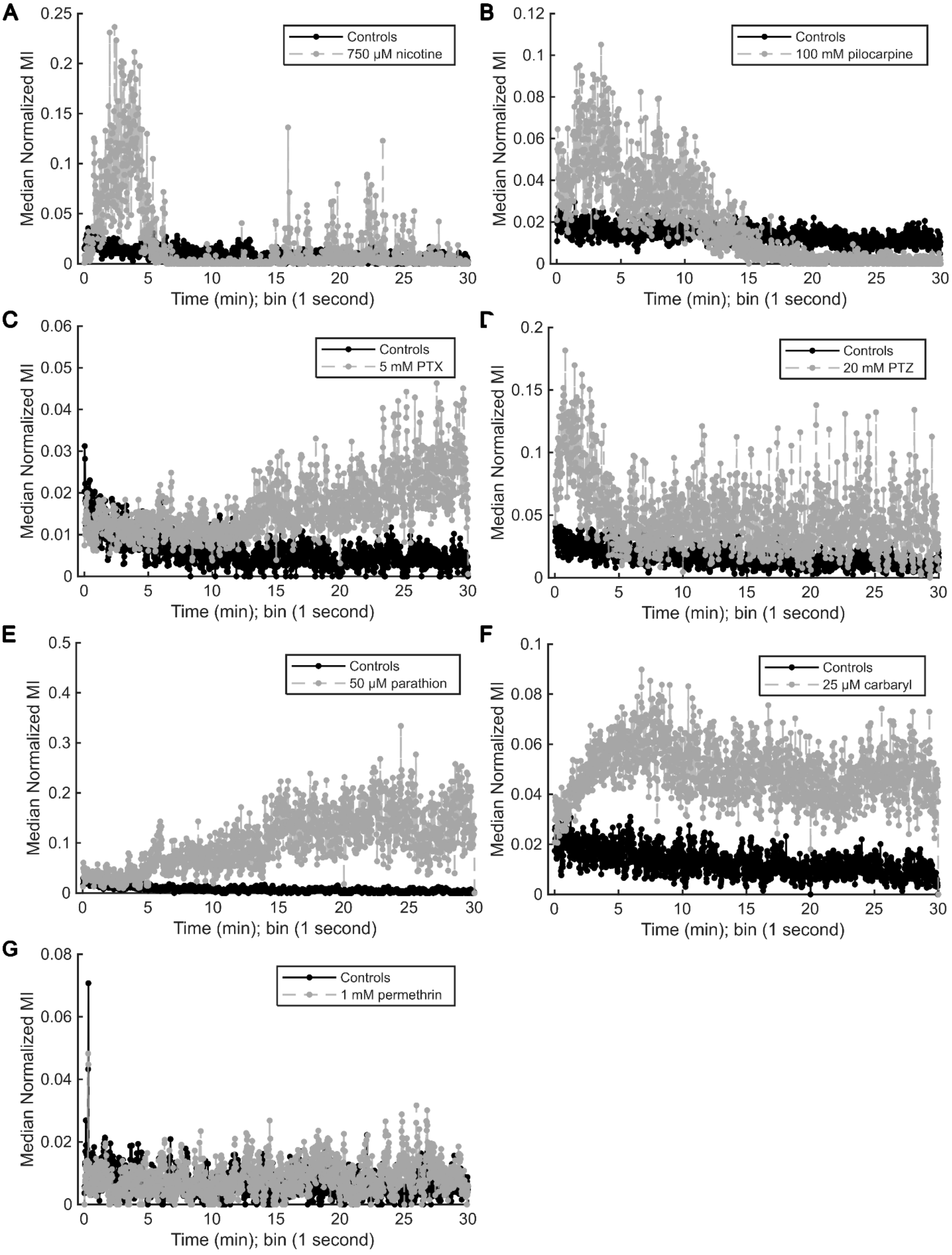
A
Controls
750 µM nicotine
Median Normalized MI
Time (min); bin (1 second)
B
Controls
100 mM pilocarpine
C
Controls
5 mM PTX
D
Controls
20 mM PTZ
E
Controls
50 µM parathion
F
Controls
25 µM carbaryl
G
Controls
1 mM permethrin

**Supplemental Figure S11. MI versus time plots for select concentrations of chemicals in DJ planarians.** The median MI of all tested planarians for each 1 second bin are shown for the concentrations with the highest number of median pSLA for each chemical compared to its in-plate vehicle control. Note the y-axes are scaled differently in the various plots to best reflect each data set. Sample sizes are listed in Supplemental Table S3.

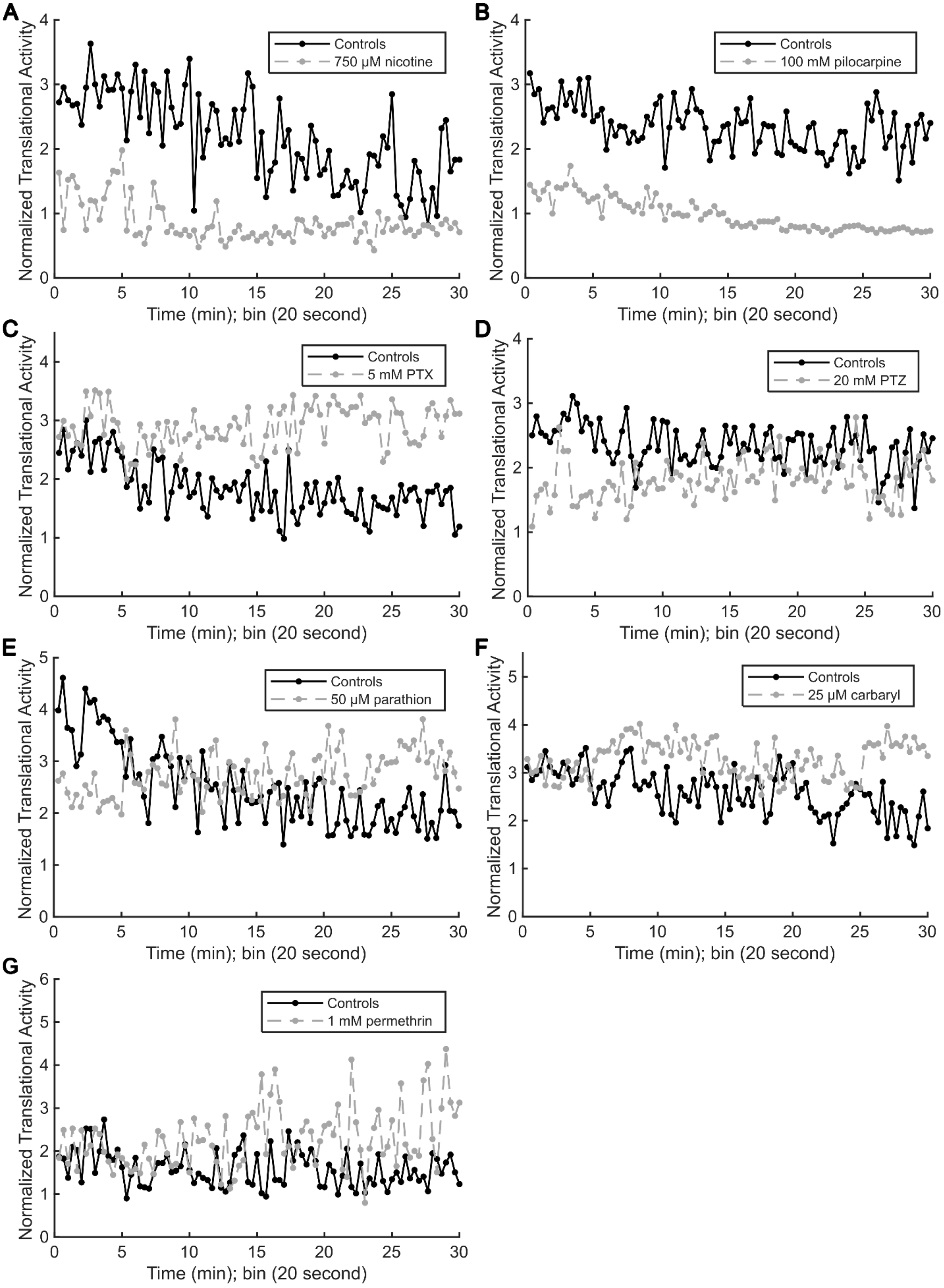
A
Controls
750 µM nicotine
B
Controls
100 mM pilocarpine
C
Controls
5 mM PTX
D
Controls
20 mM PTZ
E
Controls
50 µM parathion
F
Controls
25 µM carbaryl
G
Controls
1 mM permethrin
Normalized Translational Activity
Time (min); bin (20 second)

**Supplemental Figure S12. Translational activity vs time for select concentrations of chemicals in DJ planarians.** The median normalized translational activity of all tested planarians for each 1 second bin are shown for the concentrations with the highest number of median pSLA for each chemical compared to its in-plate vehicle control. Note the y-axes are scaled differently in the various plots to best reflect each data set. Sample sizes are listed in Supplemental Table S3.

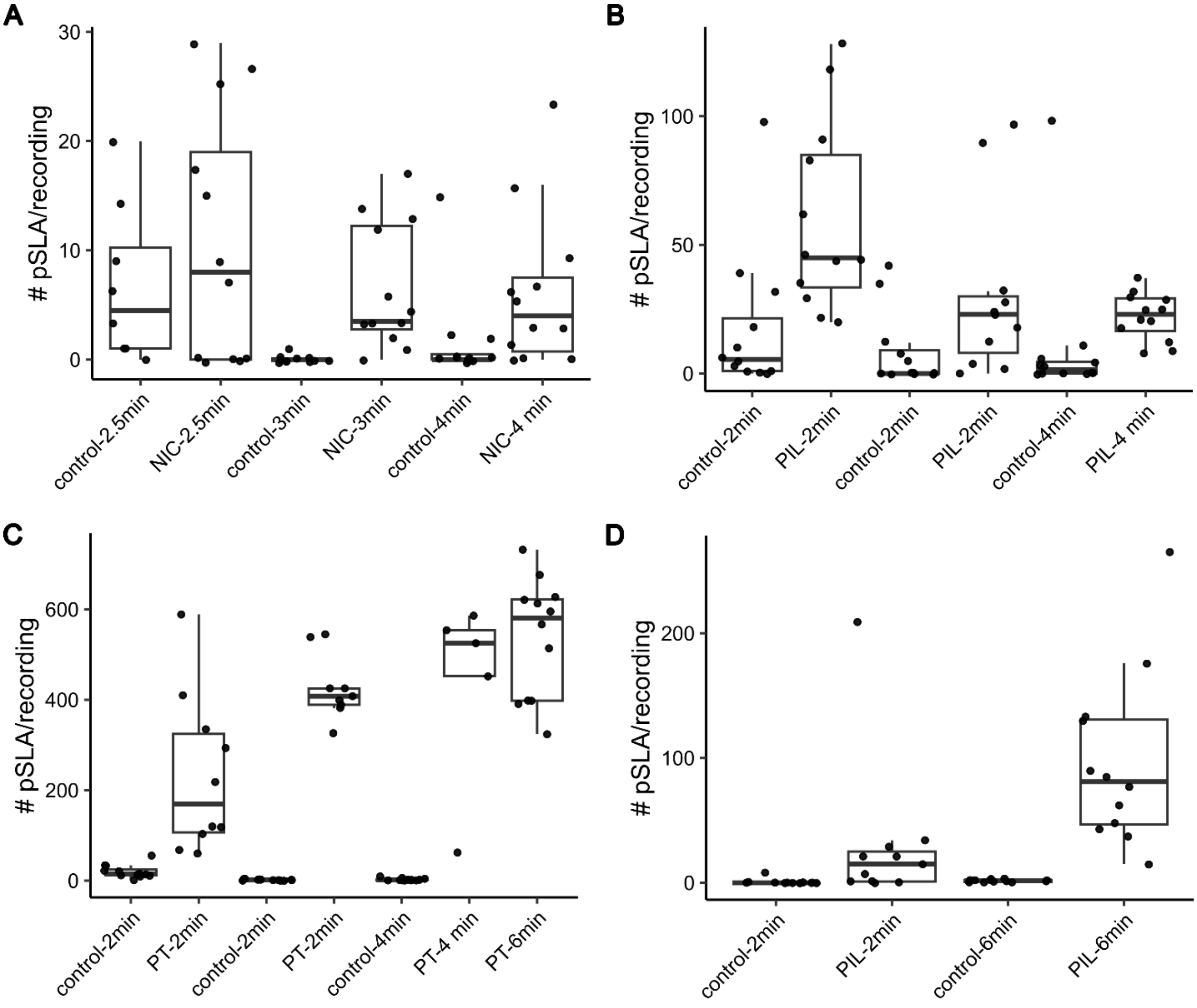


**Supplemental Figure S13. Differences in start time of recording within 2-6 minutes of chemical exposure do not drastically change results.** Boxplots show the number of pSLA events for A) GD planarians exposed to 10 µM nicotine (NIC, first 5 minutes of recording only), B) GD planarians exposed to 6 mM pilocarpine (PIL), C) DJ planarians exposed to 50 µM parathion (PT), and D) 100 mM PIL. Data are shown for each tested plate separately and list the starting time of recording as the number of minutes after chemical exposure. Each pair of control and condition represent one plate. No control is shown for "PT-6 min" in D as no adult controls were in this plate. While there is plate to plate variability in the number of pSLA events, differing delay times before starting recording do not dramatically change the results. N=12 per condition.

### Supplemental Tables

#### Supplemental Table S1. pH of highest test concentrations of each chemical.

| Chemical | Highest tested concentration in planarian water | pH |
|---|---|---|
| allyl isothiocyanate | 100 µM | 7.74 |
| carbaryl | 100 µM (0.5% DMSO) | 7.65 |
| dimethyl sulfoxide | 5% (v/v) | 7.90 |
| N-methyl-D-aspartate | 3 mM (GD) | 3.57 |
| N-methyl-D-aspartate | 15 mM (DJ) | 2.97 |
| (−)-nicotine | 1 mM | 8.75 |
| parathion | 100 µM[1](0.5% DMSO) | 7.99 |
| pentylenetetrazole | 25 mM | 6.25 |
| permethrin (mixture of cis and trans isomers) | 1 mM[1] (0.5% DMSO) | 7.95 |
| picrotoxin, *Anamirta cocculin* | 5 mM[1] | 7.25 |
| (+)-pilocarpine hydrochloride | 100 mM | 5.09 |

[1] indicates solubility issues at this concentration

#### Supplemental Table S2. Components used in imaging set up.

| Item | Ordering Info / Specifications/Cat # | Vendor | Comment |
|---|---|---|---|
| 48-well plate | 25-108MP | Genesee Scientific | Tissue culture treated |
| Camera | FL3-U3-13E4MC | Teledyne FLIR | Other camera with similar properties can be used |
| computer | Intel i5-10505 CPU @3.20 GHz, 32GB RAM; Windows 11Pro | DELL | Other computer with similar specifications can be used |
| Fresnel lens | 8.3" x 11.75" LARGE 3X Fresnel Lens FULL PAGE Magnifier | Amazon | Cut to fit the size of the 48-well plate |
| LED light panel | Tracer A4 LED Light Box 9x12 Inch | Amazon | Powered by USB; 3 light intensities |
| Lens for camera | 1:14 25 mm | Tamron | |
| Lux meter | Dr.meter LX1330B | Amazon | Reads in range of 0-200,000 lux |
| Recording box | NEEWER Photo Studio Shooting tent | Amazon | 16 inches/40cm Shooting Tent – other models work, too |
| Ring stand with clamps | 80-90 cm tall | Fisher Scientific | To hold cameras; any work |
| Thermal sealing film | TSS-RTQ-100 | Excel Scientific | For safe recording of chemicals outside the fume hood |
| USB 3 cable | CY Micro USB 3.0 Dual Screws Locking to Type-A USB 3.0 Data 5Gbps Power Cable 90 Degree Down Angled Type | Amazon | To connect camera to PC |

**Supplemental Table S3. Sample sizes associated with each figure and table.** While all conditions were tested in at n≥12, some sample sizes are lower due to worms crawling out of the well during recording and being excluded from analysis. NIC: nicotine, PIL: Pilocarpine; CAR: carbaryl, PT: parathion

| Figure or Table | N's |
|---|---|
| Figure 4A | Controls:35, NMDA: 28, pH: 12 |
| Figure 4B | Controls:24, NMDA: 19, pH: 14 |
| Figure 4C, E | Controls:35, NMDA: 28 |
| Figure 4D, F | Controls:24, NMDA: 19 |
| Figure 5A,C, E | Controls: 36, AITC: 21 |
| Figure 5B, D, F | Controls: 12, AITC: 31 |
| Figure 6A | Controls: 32, 10: 36, 15: 24, 25: 24, 50: 12 |
| Figure 6B | Controls: 24, 50: 12, 100: 12, 500: 12, 750: 12, 1000: 24 |
| Figure 6C | Controls:48 , 2: 12, 3: 12, 4: 12, 6: 35 |
| Figure 6D | Controls: 72, 6: 12, 20: 24, 40: 20, 80: 12, 100: 23 |
| Figure 6E | Controls: 36, PTX: 35 |
| Figure 6F | Controls: 24, PTX: 24 |
| Figure 7A | Controls: 24, 5: 12, 10: 12, 15:12, 20:12 |
| Figure 7B | Controls: 24, 5: 12, 10: 12, 15:24, 20:12 |
| Figure 7C | Controls: 44, 10: 12, 25: 11, 31.6: 34, 50: 22 |
| Figure 7D | Controls: 36, 10: 12, 25: 12, 50: 24, 75: 21 |
| Figure 7E | Controls: 23, 10: 21, 25: 11, 50: 21, 75: 10 |
| Figure 7F | Controls: 36, 10: 12, 25: 24, 50: 12, 75: 12 |
| Figure 7G | Controls: 24, 100: 12, 1000: 12 |
| Figure 7H | Controls: 24, 100: 12, 1000: 12 |
| Table 3; Adults | NMDA:19, pH: 14, NIC: 12, PIL:23, PTX: 24, PTZ:12; PT:24, CAR:24 |
| Table 3; Regenerating | NMDA:17, pH: 18, NIC: 12, PIL: 9, PTX: 12, PTZ:12; PT:12, CAR: 12 |
| Supplemental Figure S1 | Control-6d: 12, PT-6d: 10, Control-6d: 12, PT-6d: 5, Control-9d: 12, PT-9d:9 |
| Supplemental Figure S2A | 0: 12, 0.1: 24, 0.5: 35, 1: 12, 2:12, 4: 8, 5: 8 |
| Supplemental Figure S2B | 0: 24, 0.5: 24, 1: 12, 2:12, 4: 12, 5: 11 |
| Supplemental Figure S6; GD | Controls: 48, NMDA:12, pH: 12, NIC: 24, PIL:12, PTX: 12, PTZ:12; PT:24, CAR:12 |
| Supplemental Figure S6; DJ | Controls: 48, NMDA:12, pH: 12, NIC: 12, PIL:12, PTX: 12, PTZ:12; PT:24, CAR:12 |
| Supplemental Figure S7A, S8A | Controls: 32, nicotine: 36 |
| Supplemental Figure S7B, S8B | Controls: 12, pilocarpine: 12 |
| Supplemental Figure S7C, S8C | Controls: 36, PTX: 35 |
| Supplemental Figure S7D, S8D | Controls: 12; PTZ: 12 |
| Supplemental Figure S7E, S8E | Controls: 12, permethrin: 12 |
| Supplemental Figure S7F, S8F | Controls: 32; parathion: 34 |
| Supplemental Figure S7G, S8G | Controls: 12, carbaryl: 12 |
| Supplemental Figure S9 | Controls: 32, 10: 36, 15: 24, 25: 24 |
| Supplemental Figure S10A | Controls: 12, nicotine: 12 |
| Supplemental Figure S10B | Controls: 11, carbaryl: 10 |
| Supplemental Figure S11A, S12A | Controls: 12, nicotine: 12 |
| Supplemental Figure S11B, S12B | Controls: 24, pilocarpine: 24 |
| Supplemental Figure S11C, S12C | Controls: 24, PTX: 24 |
| Supplemental Figure S11D, S12D | Controls: 12, PTZ: 12 |
| Supplemental Figure S11E, S12E | Controls: 12, permethrin: 12 |
| Supplemental Figure S11F, S12F | Controls: 24; parathion: 24 |
| Supplemental Figure S11G, S12G | Controls: 24, carbaryl: 24 |

**Supplemental Table S4. Lowest concentration to statistically significantly induce pSLA in each species.** N/A indicates none of the tested concentrations induced pSLA.

| Chemical | GD | DJ |
|---|---|---|
| NMDA | 15 mM | 3 mM |
| Nicotine | N/A (10 µM, first 5 min) | 100 µM |
| Pilocarpine | 2 mM* | 40 mM |
| PTX | 5 mM* | 5 mM* |
| PTZ | 15 mM | 10 mM |
| Parathion | 10 µM | 10 µM |
| Carbaryl | N/A | 25 µM |
| Permethrin | N/A | N/A |

*lowest concentration tested

**Supplemental Table S5**. **Median number of pSLA in GD planarians exposed to representative concentrations of seizurogenic compounds.** Sample sizes are listed in Supplemental Table S3 for the associated figure.

| Condition | Median # pSLA (25th, 75th percentile) | Associated figure |
|---|---|---|
| 3 mM NMDA | 84 (57, 133)* | 4A |
| pH 3.65 | 14 (7, 35)* | 4A |
| 10 µM nicotine | 6 (2, 17) | 6A |
| 3 mM pilocarpine | 48 (21, 79)* | 6C |
| 5 mM PTX | 16 (5, 31)* | 6E |
| 15 mM PTZ | 22 (5,28)* | 7A |
| 31.6 µM parathion | 89 (53, 148)* | 7C |
| 25 µM carbaryl | 18 (8, 25) | 7E |

* significantly different from controls ($p<0.05$) using pairwise contrasts (with a Benjamini & Hochberg p-value correction) of the estimated marginal means of a negative binomial generalized linear model of condition versus number of pSLA events

**Supplemental Table S6**. **Post-hoc power analysis for statistically significant conditions.** The achieved power was calculated in R by running simulated data based on the parameters (group means, sample size and theta calculated from the negative binomial generalized linear model) through the same statistical analysis pipeline. Focusing on pairwise concentrations with the controls that showed statistical significance in our original analysis, power was calculated as the number of times that specific pairwise comparison was simulated to be significant ($p<0.05$) divided by the number of simulations (n=5000). Power was heavily influenced by theta, a measure of dispersion, which was much lower in GD planarians, demonstrating greater data dispersion.

| Species | Associated figure | Tested conditions | Theta | Power for statistically significant conditions |
|---|---|---|---|---|
| GD | 4A | 3 mM NMDA*, pH 3.6* | 0.85 | 1, 0.43 |
| GD | 6C | 2*, 3*, 4*, 6* mM pilocarpine | 0.64 | 0.86, 0.94, 0.9, 0.98 |
| GD | 6E | 5 mM PTX* | 0.42 | 0.53 |
| GD | 7A | 5, 10, 15, 20 mM PTZ | 0.58 | 0.59, 0.6642 |
| GD | 7C | 10*, 25*, 31.6*, 50* µM PT | 0.98 | 0.974, 1, 1, 1 |
| DJ | 4B | 15 mM NMDA*, pH 3* | 0.77 | 1, 1 |
| DJ | 6B | 50, 100*, 500*, 750*, 1000* µM nicotine | 1.55 | 0.59, 1, 1, 1 |
| DJ | 6D | 6, 20, 40*, 80*, 100* mM pilocarpine | 0.6 | 0.93, 1, 1 |
| DJ | 6F | 5 mM PTX | 2.1 | 1 |
| DJ | 7B | 5, 10*, 15*, 20* mM PTZ | 1.86 | 1, 1, 1 |
| DJ | 7D | 10*, 25*, 50*, 75* µM PT | 1.67 | 1, 1, 1, 1 |
| DJ | 7F | 10, 25*, 50, 75 µM carbaryl | 0.65 | 0.87 |

* significantly different from controls ($p<0.05$) using pairwise contrasts (with a Benjamini & Hochberg p-value correction) of the estimated marginal means of a negative binomial generalized linear model of condition versus number of pSLA events

#### Supplemental Videos

**Supplemental Video S1. Example of pSLA behavior in a GD planarian exposed to 3 mM NMDA.** Scale bar: 1 mm. Video is shown in real time (imaged at 5 frames per second).

**Supplemental Video S1. Example of pSLA behavior in a DJ planarian exposed to 3 mM NMDA.** Scale bar: 1 mm. Video is shown in real time (imaged at 5 frames per second).

#### Supplemental File Legends

**Supplemental File S1. Raw pSLA data and statistics for every condition.** NaNs represent planarians that were excluded from analysis due to crawling out of the solution during recording or were added to allow for matrices of the same size when sample sizes differed across conditions. Medians and 25th and 75th percentiles for each condition are listed below the pSLA counts. P-values for pairwise comparisons of the estimated marginal means of a negative binomial generalized linear model are listed to the right of each comparison.
